# HIRA-mediated H3.3 deposition preserves hepatocyte cell identity during liver aging

**DOI:** 10.64898/2026.03.09.710643

**Authors:** Rouven Arnold, Marcos Garcia Teneche, Xue Lei, Armin Gandhi, Christina Huan Shi, Jessica Proulx, Adarsh Rajesh, Aaron P. Havas, Shiqi Su, Annanya Sethiya, Shanshan Yin, Hiroshi Tanaka, Zong-Ming Chua, Andrew Davis, Laurence Haddadin, Michael Alcaraz, Irene Huang, Angela Liou, Anais Equey, Nirmalya Dasgupta, Karl N. Miller, Margaret Tulessin, Buddy Charbono, Adriana Charbono, Siva Karthik Varanasi, Rebecca A. Porritt, Guillermina Garcia, Sakshi Chauhan, Brian Egan, Michael Choob, Carolin Mogler, Kevin Y. Yip, Keiko Ozato, Susan M. Kaech, Yu Xin Wang, Peter D. Adams

## Abstract

Age-associated functional decline is partly driven by progressive chromatin degeneration. Maintenance of chromatin integrity preserves cell identity and promotes healthy aging, but through different mechanisms in proliferating and non-proliferating cells. However, specific mechanisms of chromatin maintenance and their compensatory capacity in proliferating and non-proliferating cells are undefined. The histone chaperone HIRA deposits the histone variant H3.3 in a DNA replication-independent manner, leading to its accumulation in aging, non-proliferating cells. Here, we show that hepatocyte-specific loss of HIRA causes loss of cell identity, metabolic dysfunction, and accelerated fibrotic pathology with age. Transcriptomic and epigenomic analyses indicate that HIRA-H3.3 preserves chromatin integrity and sustains transcription of highly expressed genes, including cell identity genes. Partial hepatectomy, associated with induced proliferation, restores identity of HIRA knockout livers with compensatory deposition of canonical histones H3.1/2. Together, these results demonstrate that HIRA-mediated H3.3 deposition is essential for safeguarding cell identity and tissue function during aging of non-proliferating cells, but this function can be rescued by tissue regeneration and associated cell proliferation.

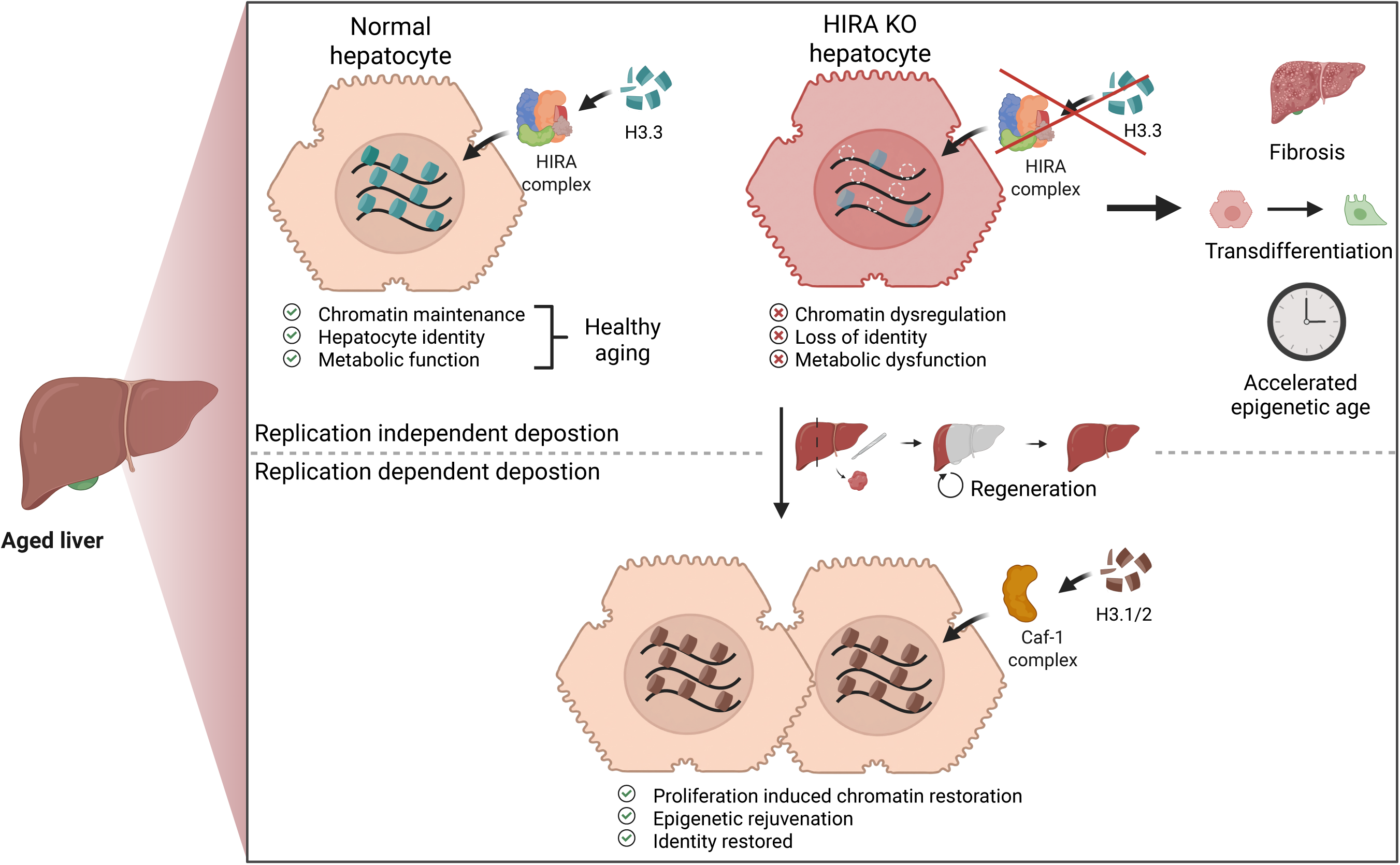

## Introduction

The epigenetic landscape governs gene expression programs and defines cell identity^1,2^. However, aging progressively erodes cell-type-specific epigenetic and transcriptional programs, driving tissue dysfunction and contributing to age-related disease and increased mortality^3–7^. Therefore, preserving cell identity by maintaining epigenetic information is vital for optimal cellular function across the lifespan.

The formation and preservation of the epigenetic landscape occurs, among other processes, through assembly, recycling, and turnover of nucleosomes, the minimum repeating units of chromatin^8^. Histones, the building blocks of nucleosomes, are predominantly deposited via two distinct mechanisms: First, replication-dependent deposition of histones and assembly of nucleosomes occurs during S-phase of the cell cycle, through the heterotrimeric CAF-1 histone chaperone complex; and second, replication-independent deposition of histone variants and nucleosome assembly takes place in non-proliferating and post-mitotic cells, through chaperones such as HIRA and ATRX/DAXX^9,10^.

ATRX/DAXX deposits H3.3 mostly at heterochromatin regions, including pericentromeres, retrotransposons, and subtelomeric regions^11^. In contrast, the HIRA histone chaperone complex, which includes HIRA, UBN1, and CABIN1, and functions together with ASF1a, deposits H3.3-H4 heterotetramers in a DNA replication-independent manner, mainly at active promoters, enhancers, and gene bodies^11–15^. HIRA and H3.3 have been implicated in development and differentiation, as well as in cellular senescence, an irreversible stress-induced growth arrest that promotes tissue aging^16–19^. Mechanistically, loss of HIRA-dependent H3.3 deposition reduces activating histone marks (H3K27ac) at lineage-specific loci while also derepressing polycomb-silenced genes^20,21^. Constitutive whole-body knockout of HIRA in mice results in gastrulation defects and embryonic lethality by midgestation^16^. Early-life lineage-specific constitutive HIRA deletion impairs cell fate commitment and tissue homeostasis across multiple cell types, causing loss of identity-defining transcription factors and ectopic activation of alternative lineage programs in skeletal muscle^20^, melanoblasts^22^, and hematopoietic stem cells^23^, as well as progressive tissue damage in cardiomyocytes^24,25^.

Surprisingly, given the role of HIRA in deposition of H3.3 at active promoters, enhancers, and gene bodies, to date no studies have specifically investigated whether HIRA is required for healthy aging of adult tissues. After development, many tissues are largely non-proliferative and therefore rely on DNA replication-independent histone assembly. H3.3 accumulates to near-saturated levels in these tissues with age, as replication-dependent variants H3.1 and H3.2 are progressively replaced by H3.3 in non-proliferative cells by ATRX/DAXX and HIRA^26–30^. The liver is one such lowly proliferative organ in which H3.3 accumulates during aging^26^. In human metabolic dysfunction-associated steatotic liver disease (MASLD) and other models of chronic liver damage, loss of liver zonation and acquisition of biphenotypic gene programs accompany disease progression^31,32^, underscoring how failure to maintain cell identity in aging and stress drives pathology in liver.

In this study, we set out to investigate the requirement for HIRA-mediated deposition of histone H3.3 for healthy aging of mouse liver hepatocytes *in vivo*. To do this, we ablated HIRA in young adult mouse liver hepatocytes at 6 months of age and aged mice to 18 months of age. HIRA knockout (KO) resulted in progressive tissue dysfunction and pathology, accompanied by specific epigenetic and transcriptomic defects and loss of cell identity. Defects were recapitulated at the molecular level by H3.3 knockout. Partial hepatectomy, associated with induced proliferation, restored cell identity. Together, these results demonstrate that HIRA-mediated H3.3 deposition is essential for safeguarding cell identity and tissue function during aging of non-proliferating cells, but this function can be rescued by tissue regeneration associated cell proliferation.

## Results

### Hepatocyte-specific HIRA deletion results in fibrosis, metabolic dysregulation, and an altered transcriptome, particularly affecting highly expressed HNF4α target genes

To investigate the requirement of HIRA for liver aging, we first confirmed histone H3.3 accumulation during normal mouse liver aging by western blot analysis of liver lysates from 4, 12, and 22-month-old female mice (Fig. 1A). Abundance of H3.3, compared to total H3 (H3.1, H3.2 and H3.3) progressively increased with age, as reported previously^26^. To probe the functional role of HIRA, we generated hepatocyte-specific HIRA KO mice by retro-orbital injection of hepatocyte-specific AAV8-Tbg-Cre into HIRA^fl/fl^ C57BL/6 mice^33^. Specifically, we injected 6-month-old male and female HIRA^fl/fl^ mice with AAV8-Tbg-Cre and aged them for an additional 12 months (Fig. 1B). The mice were closely monitored over these 12 months and showed no significant changes in body weight, abdominal swelling suggestive of neoplasia, nor other signs of malaise (Suppl. Fig. 1A). At the end of this time, the mice were culled and tissues and plasma harvested for analysis (Fig. 1B). Western blot analysis of whole liver lysates from both male and female HIRA^fl/fl^ AAV8-Tbg-Cre mice confirmed efficient HIRA KO compared to controls (wildtype injected with AAV8-Tbg-Cre and HIRA^fl/fl^ injected with AAV8-Tbg-MCS (AAV ctl)) (Fig. 1C, Suppl. Fig. 1B). Immunofluorescence staining validated reduced H3.3 deposition in euchromatin, while perinuclear staining persisted, consistent with continued DAXX-mediated H3.3 deposition in repetitive heterochromatin (Suppl. Fig. 1C). Plasma liver damage markers (AST, ALP) showed no significant change in mice with HIRA KO liver (Suppl. Fig. 1D-E). To assess liver histology, we performed hematoxylin and eosin (H&E) and picrosirius red staining on liver sections. Quantification of picrosirius red staining revealed a significant increase in liver fibrosis in HIRA KO mice compared to controls (Fig. 1D, E).

**Figure 1:**
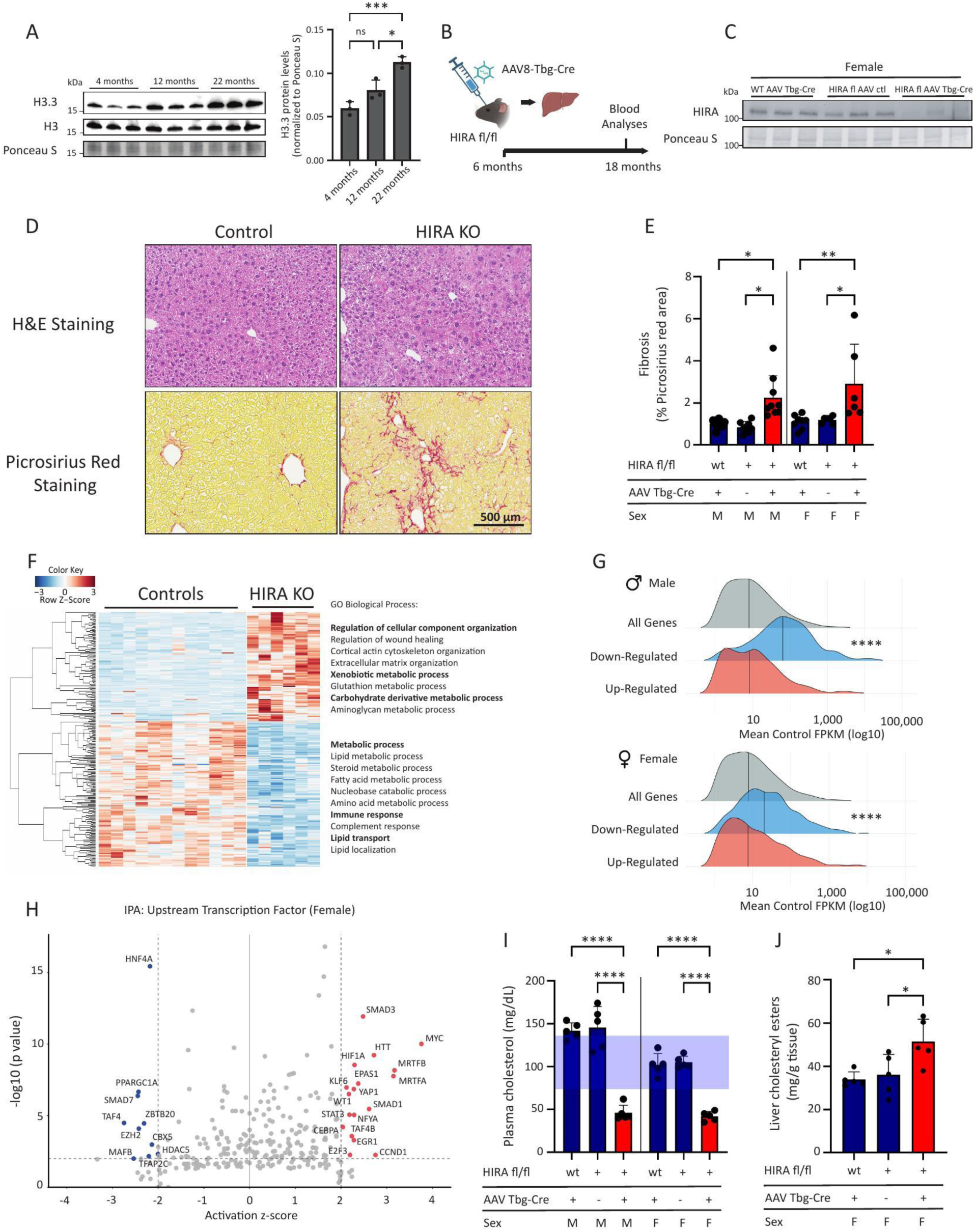
Hepatocyte-specific HIRA deletion results in fibrosis, metabolic dysregulation, and an altered transcriptome, particularly affecting highly expressed HNF4α target genes. A. Western blot analysis of HIRA, H3.3, and total H3 of liver lysates from ∼4, 12, and 22-month-old female mice, with quantification. Ponceau S staining was used as loading control. n = 3 per age group. B. Graphical representation of the experimental design for hepatocyte-specific HIRA deletion. C. Representative western blot demonstrating HIRA depletion in liver lysates from control and HIRA KO female mice. Ponceau S staining was used as loading control. n = 8 (WT AAV Tbg-Cre); n = 6 (HIRA^fl/fl^ AAV ctl, HIRA^fl/fl^ AAV Tbg-Cre). D. Representative hematoxylin and eosin (H&E) and picrosirius red stainings of liver section from control and HIRA KO mice. Scale bar, 500 μm. E. Quantification of picrosirius red positive area. Male mice: n = 9 (WT AAV Tbg-Cre, HIRA^fl/fl^ AAV Tbg-Cre); n = 8 (HIRA^fl/fl^ AAV ctl), Female mice: n = 8 (WT AAV Tbg-Cre); n = 6 (HIRA^fl/fl^ AAV ctl, HIRA^fl/fl^ AAV Tbg-Cre). F. Heatmap of RNA-seq data comparing male and female controls (WT AAV Tbg-Cre, HIRA^fl/fl^ AAV ctl) with HIRA KO (HIRA^fl/fl^ AAV Tbg-Cre) mice. n = 3 per group. G. Ridgeline plot illustrating expression distribution of all expressed genes compared to significantly up- and down-regulated genes. H. Ingenuity Pathway Analysis (IPA) showing top predicted upstream transcription factors regulating differentially expressed genes (DEGs) in control versus HIRA KO mice. I. Plasma cholesterol levels in control and HIRA KO mice. n = 5. J. Liver choleteryl esters normalized to tissue weight from control and HIRA KO mice. n = 5. Data are presented as means ± SEM. Statistical significance was determined by one-way analysis of variance (ANOVA) with Tukey’s multiple comparisons test, except for (G), where the Wilcoxon rank-sum test (Mann-Whitney U test) was used. Asterisk (*) indicates p < 0.05, (**) p < 0.01, (***) p< 0.001, (****) p < 0.0001.

To understand the molecular basis of this phenotype, we performed RNA-seq on liver tissue from control and HIRA KO. Principal component analysis (PCA) analysis showed clear separation among the groups, with only minor sex-dimorphic effects (Suppl. Fig. 1F). Differential expression analysis identified 145 genes that were significantly downregulated, while 107 genes were found to be significantly upregulated in male and female HIRA KO livers (Fig. 1F). Gene ontology analysis of upregulated genes showed enrichment for processes associated with the fibrotic phenotype, including ‘regulation of cellular component organization’, ‘regulation of wound healing’, ‘extracellular matrix organization’, and ‘xenobiotic metabolic process’. In contrast, the majority of downregulated genes were associated with metabolic processes, including ‘lipid metabolic process’, ‘lipid transport’ and ‘amino acid metabolic process’, suggesting impaired hepatic metabolic function (Fig. 1F). Notably, downregulated genes were significantly enriched for highly expressed liver genes in both male and female mice, distinguishing them from the overall transcriptome and from upregulated genes (Fig. 1G). To identify transcriptional regulators driving these gene expression changes, we performed Ingenuity Pathway Analysis (IPA). This analysis predicted inhibition of HNF4α, a master regulator of hepatocyte identity and metabolism, in both sexes. Additional metabolic transcription factors showed sex-specific inhibition patterns, including SREBF1 and SREBF2 in males and PPARGC1A and ZBTB20 in females (Fig. 1H, Suppl. Fig. 1G). Given the predicted HNF4α inhibition and enrichment of lipid metabolic processes among downregulated genes, we examined cholesterol homeostasis, a critical hepatic lipid pathway. In the liver, cholesterol is synthesized through a multi-step pathway from acetyl-CoA and can then be converted to bile acids or packaged into lipoprotein particles for export^34^. Strikingly, examination of cholesterol distribution showed opposing patterns between plasma and liver: total plasma cholesterol was significantly decreased in HIRA KO male and female mice (Fig. 1I), while lipidomic analysis of female liver samples revealed significantly increased cholesteryl ester content, with specific accumulation of cholesteryl oleate (18:1) and cholesteryl linoleate (18:2), the predominant esterified cholesterol species in mammalian liver (Fig. 1J, Suppl. Fig. 1H). This inverse relationship suggests impaired cholesterol export from the liver. Consistent with this, while plasma lipoprotein analysis showed that HDL, LDL, and VLDL cholesterol levels were all significantly reduced in male and female HIRA KO mice (Suppl. Fig. 1I), analysis of plasma bile acids (synthesized in the liver from cholesterol) revealed a significant increase in HIRA KO male mice and a marked upward trend in female mice (Suppl. Fig. 1J). These findings demonstrate that KO of HIRA in hepatocytes results in fibrosis, metabolic dysregulation, and an altered transcriptome, particularly affecting highly expressed HNF4α target genes.

### HIRA loss causes erosion of hepatocyte identity and trans-differentiation

To better understand how loss of HIRA activity affects hepatocytes, we performed a time-course analysis to assess maintenance of tissue and cell identity. To do this, we employed an alternative method of Adeno-associated virus (AAV) CRISPR-mediated hepatocyte-specific HIRA KO, using AAV8-Tbg-saCAS9/U6-sgRNA-HIRA or AAV8-Tbg-saCAS9/U6-sgRNA-Rosa26 as control^35^, in male mice for analysis at 3 weeks and 6 months after KO. Western blot analysis comparing CRISPR-mediated HIRA KO to Rosa26-targeted control mice confirmed loss of HIRA protein expression (Suppl. Fig. 2A). We performed bulk RNA sequencing of HIRA KO livers at 3 weeks and 6 months. PCA indicated that the control and KO groups are distinctly separated (Suppl. Fig. 2B). Differential expression analysis identified 290 upregulated and 145 downregulated genes at 3 weeks, and 227 upregulated and 373 downregulated genes at 6 months after KO. Circular visualization of DEGs and associated GO terms at all three timepoints (3 weeks, 6 months, and 1 year) is shown in Fig. 2A, B. Gene Ontology analysis revealed strikingly different temporal patterns: upregulated genes revealed that stress-associated pathways, including ‘extracellular matrix organization’, ‘wound response’, and ‘growth factor signaling’, were largely absent at early timepoints and predominantly emerged at 1 year (Fig. 2A, bottom). In contrast, downregulated genes showed consistent GO term enrichment across all three timepoints, with hepatocyte-specific metabolic functions including ‘regulation of lipid metabolic process’, ‘amino acid metabolism’, ‘fatty acid metabolism’, ‘steroid metabolic process’, and ‘bile secretion’ downregulated as early as 3 weeks and persisting through 1 year (Fig. 2B, bottom). Consistent with the delayed emergence of stress-associated pathways, fibrosis levels showed no significant differences between control and HIRA KO livers at 3 weeks or 6 months (Suppl. Fig. 2C, D). Given the consistent downregulation of metabolic processes, we quantified ‘liver identity’ using Gene Set Variation Analysis (GSVA) with a liver-specific gene signature comprising 85 genes^36^. Liver identity scores were significantly reduced at 3 weeks, 6 months, and 1 year post-KO compared to age-matched controls (Fig. 2C). Taken together, these results suggest that the stress-associated fibrotic response is a secondary delayed effect, while the loss of identity is a primary consequence of HIRA KO.

**Figure 2:**
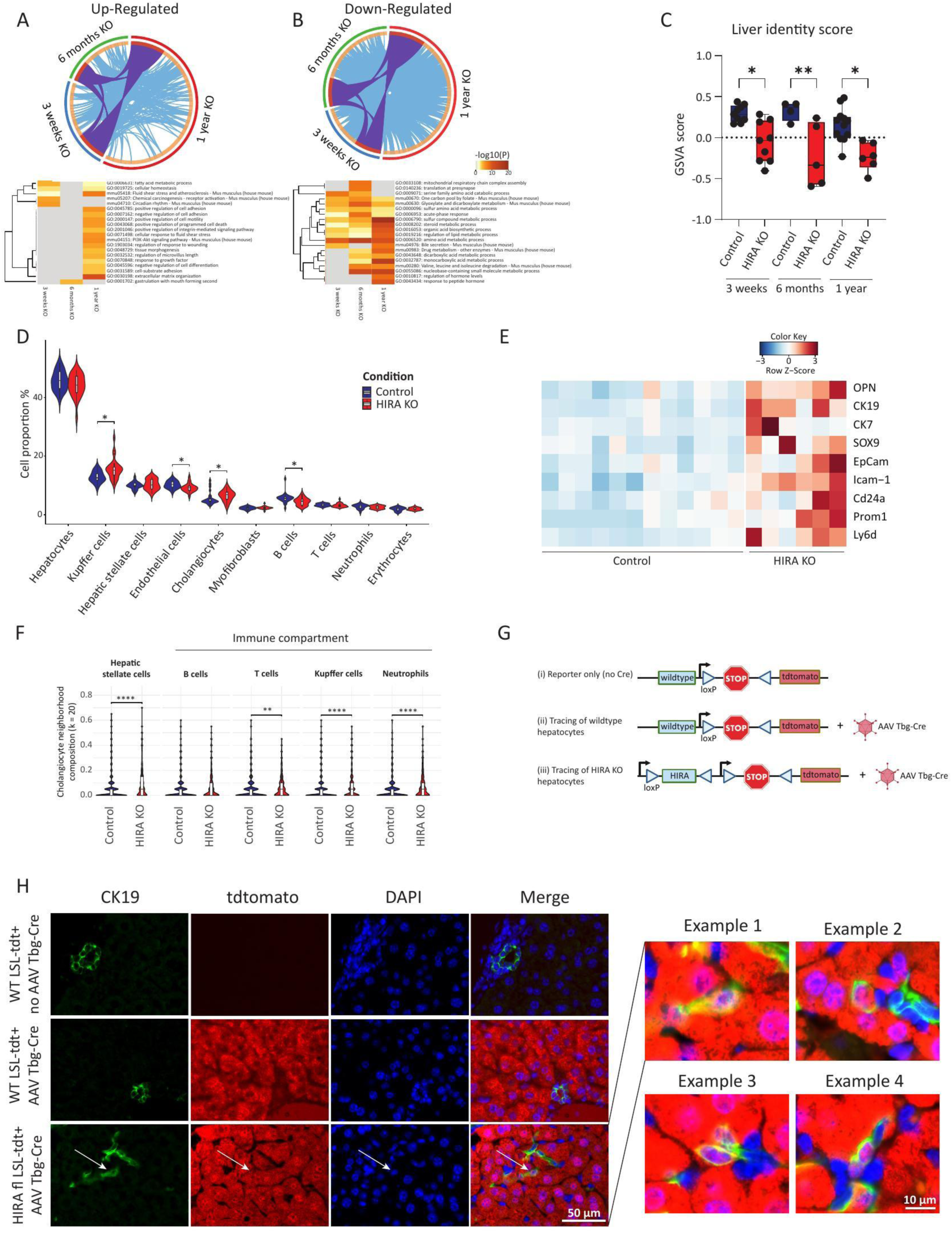
HIRA loss causes erosion of hepatocyte identity and trans-differentiation. A, B. Metascape analysis reveals gene overlap and shared gene ontology (GO) terms among DEGs from female HIRA KO datasets at 3 weeks (CRISPR), 6 months (CRISPR), and 1 year (Cre). Purple curves connect identical genes across datasets; blue curves connect genes enriched within the same ontology term. n = 5 (3-week and 6-month KO); n = 3 (1-year KO). C. Gene set variation analysis (GSVA) of 85 liver identity genes in control and HIRA KO mice across time points. n = 5 (3-week and 6-month KO); n = 6 (1-year KO). D. Relative abundance of cell populations in control (WT AAV Tbg-Cre) and HIRA KO (HIRAfl/fl AAV Tbg-Cre) liver sections as determined by CosMX spatial transcriptomics analysis. n = 3 per group. E. Heatmap of classical biliary lineage marker expression from RNA-seq analysis comparing male and female controls (WT AAV Tbg-Cre, HIRA^fl/fl^ AAV ctl) with HIRA KO (HIRA^fl/fl^ AAV Tbg-Cre) mice. n = 3 per group. F. Spatial neighborhood analysis of cholangiocytes (k = 20 nearest neighbors) in control and HIRA KO liver sections, showing the relative abundance of hepatic stellate cells, B cells, T cells, Kupffer cells, and neutrophils within the cholangiocyte microenvironment. n = 3 per group. G. Schematic illustrating hepatocyte lineage tracing strategy. AAV-Tbg-Cre was used to induce hepatocyte-specific recombination of the Rosa26-LSL-tdTomato reporter, with or without HIRA deletion. H. Lineage tracing liver sections from control WT LSL-tdTomato without virus injection, WT LSL-tdTomato with AAV Tbg-Cre injection and HIRA KO HIRAfl/fl LSL-tdTomato with AAV Tbg-Cre injection male mice were analyzed by immunofluorescence microscopy. Sections were stained with DAPI, tdTomato and CK19 antibodies. Scale bar, 50 μm for the overview images, 10 μm for the individual examples. Data are presented as means ± SEM. Statistical significance for C was determined by one-way analysis of variance (ANOVA) with Tukey’s multiple comparisons test. For D and G, the Wilcoxon rank-sum test was used. Asterisk (*) indicates p < 0.05, (**) p < 0.01, (***) p< 0.001, (****) p < 0.0001.

To examine the cellular composition and spatial organization of this loss of identity phenotype at single-cell resolution, we performed CosMx spatial transcriptomics analysis on 1 year post-HIRA KO and control livers, using a 1000 cell identity gene panel (Fig. 2D, F, Suppl. Fig. 2E-G). Unsupervised clustering identified ten distinct liver cell populations including hepatocytes, cholangiocytes, hepatic stellate cells, endothelial cells, Kupffer cells, myofibroblasts, neutrophils, T cells, B cells, and erythrocytes (Suppl. Fig. 2E). Cell type assignments from spatial transcriptomics were validated by dot plot analysis examining expression of top differentially expressed marker genes (log2FC >1) across all identified populations (Suppl. Fig. 2F). Consistent with the histologically observed fibrotic phenotype (Fig. 1D), a composite fibrosis gene signature score in hepatic stellate cells revealed significant elevation in HIRA KO livers relative to controls (Suppl. Fig. 2G, Suppl. Table 1). Cell type proportion analysis demonstrated significant expansion of cholangiocyte and Kupffer cell populations in HIRA KO livers, alongside significant decrease of endothelial cell and B cell populations (Fig. 2D). This enrichment of cholangiocyte markers was confirmed by bulk RNA-seq data after 1 year of HIRA KO (Fig. 2E). Spatial neighborhood analysis (k=20 nearest neighbors) of the cholangiocyte microenvironment revealed that cholangiocytes in HIRA KO livers were surrounded by significantly elevated proportions of hepatic stellate cells, T cells, Kupffer cells, and neutrophils compared to controls (Fig. 2F), suggesting that these cells are embedded within an inflammatory and pro-fibrotic microenvironment characteristic of liver injury.

Hepatocytes can transdifferentiate to cholangiocytes in response to chronic liver injury^31,37^, so we investigated whether a hepatocyte-to-cholangiocyte conversion was occurring in HIRA KO livers. To do this, we performed lineage tracing using a fluorescent reporter by retro-orbital injection of AAV8-Tbg-Cre into HIRA^fl/fl^; LSL-tdtomato mice (Fig. 2G). Immunofluorescence co-staining for CK19 (green), tdTomato (red), and DAPI (blue) showed that in wildtype LSL-tdTomato mice that received no AAV, CK19+ cholangiocytes and other cells displayed no tdTomato signal (Fig. 2H, top row). Wildtype LSL-tdTomato mice that received AAV8-Tbg-Cre displayed robust tdTomato labeling in hepatocytes with normal CK19+ cholangiocyte abundance and distribution, and only rare tdTomato+/CK19+ double-positive cells (Fig. 2H, middle row). In contrast, HIRA^fl/fl^; LSL-tdTomato mice injected with AAV8-Tbg-Cre showed numerous tdTomato+/CK19+ double-positive cells (indicated by white arrows), confirming hepatocyte-to-cholangiocyte transdifferentiation (Fig. 2H, bottom row). High-magnification examples from four representative regions demonstrate the morphology and spatial distribution of these lineage-traced transdifferentiated cells, showing the colocalization of tdTomato and CK19 signals (Fig. 2H, right panels). Collectively, these results demonstrate that hepatocyte-specific HIRA KO leads to loss of hepatocyte identity and their transdifferentiation into cholangiocytes.

### HIRA preserves chromatin integrity and epigenetic age

Given that HIRA is a histone chaperone and the epigenome determines cell identity, we set out to assess the impact of HIRA KO on the epigenome. To do this, we performed a comprehensive epigenetic analysis including ATAC-seq, DNA methylation array, and CUT&Tag for H3.3, H3K27ac, and H3K36me3 on control and HIRA KO liver 1 year after KO. We first examined chromatin accessibility by ATAC-seq. We identified 4,740 peaks with increased accessibility and 2,312 peaks with decreased accessibility in HIRA KO (Suppl. Fig. 3A). The genomic distribution of differentially accessible peaks revealed that 81 % mapped to intronic and intergenic regions, 4 % mapped to untranslated regions (UTR), 5 % to transcription termination sites (TTS), 5 % to exons, 4 % to transcription start sites (TSS), and 0.6% to non-coding regions (Suppl. Fig. 3B), indicating that HIRA regulates chromatin structure broadly across the genome, including at regulatory regions, as previously reported^12,13,38^. Consistent with a direct regulatory relationship, changes in TSS accessibility at DEGs positively correlated with their expression changes in HIRA KO livers (Fig. 3A). In line with decreased expression of HNF4α target genes (Fig. 1H and Fig. Suppl. 1G), motif analysis revealed decreased enrichment of the HNF4α binding motif in accessible chromatin of HIRA KO samples (Suppl. Fig. 3C). To examine DNA methylation changes, we performed methylation array profiling and identified 35,594 hypomethylated and 7,968 hypermethylated CpG sites in HIRA KO liver (Suppl. Fig. 3D). Genomic annotation of differentially methylated sites showed that both hyper- and hypomethylated CpGs mapped across promoter, intronic, intergenic, and exonic regions (Suppl. Fig. 3E). Annotation of the methylation data onto the observed correlation between TSS accessibility and gene expression changes revealed a coherent epigenetic pattern: downregulated genes exhibited both reduced TSS accessibility and promoter DNA hypermethylation, whereas upregulated genes showed increased TSS accessibility and promoter hypomethylation (Fig. 3A). These coordinated changes are consistent with established principles linking increased gene expression with chromatin accessibility and DNA hypomethylation at promoters^39^. Next, using the methylation data, we applied the Zhou347 epigenetic clock, a multi-tissue age predictor, and found that HIRA KO livers displayed a significant increase in predicted biological age relative to controls (Fig. 3B)^40^. Taken together, these results suggest that HIRA KO results in integrated changes in chromatin accessibility and DNA methylation at promoter/TSSs that, as expected, correlate with changes in gene expression and are associated with increased epigenetic age.

**Figure 3:**
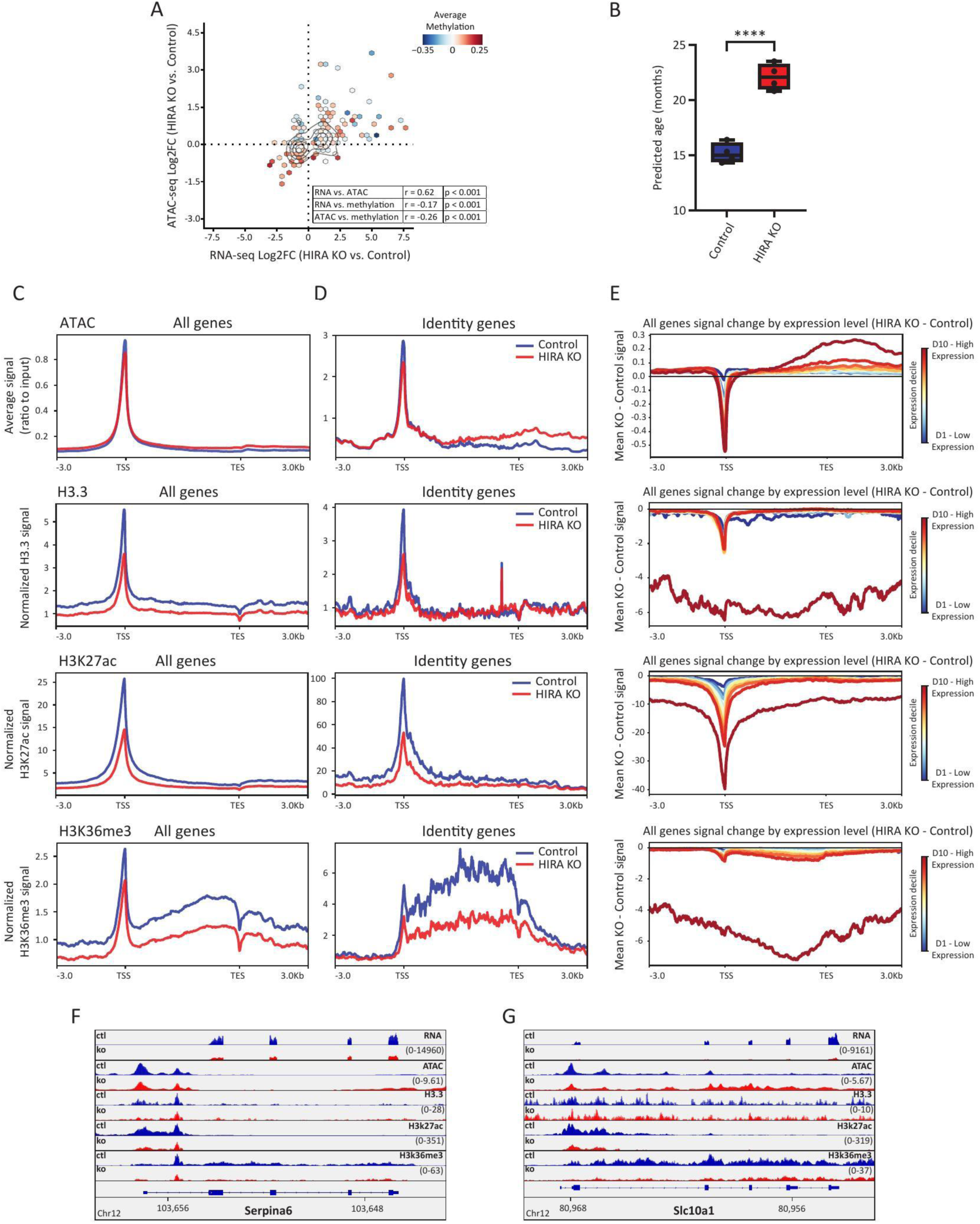
HIRA preserves chromatin integrity and epigenetic age. A. Hexbin plot of RNA-seq (log2 fold change) versus ATAC-seq (log2 fold change) for DEGs, colored by average DNA methylation change at each bin. All DEGs from the 1-year HIRA KO bulk RNA-seq dataset (Fig. 1) are plotted. n = 3 per group for the RNA-seq, n = 2 per group for the ATAC-seq, n = 4 per group for the DNA methylation array. B. Epigenetic age prediction using the Zhou347 DNA methylation-based clock in control and HIRA KO livers. n = 4 per group. C. Metaplots of ATAC-seq, H3.3, H3K27ac, and H3K36me3 CUT&Tag signal across all protein-coding genes (TSS to TES ± 3.0 Kb) in control and HIRA KO livers. Signal is shown as average ratio to input (ATAC) or normalized signal (CUT&Tag). n = 2 per group (ATAC), n = 3 per group (CUT&Tag). D. Metaplots of ATAC-seq, H3.3, H3K27ac, and H3K36me3 CUT&Tag signal at liver identity genes (TSS to TES ± 3.0 Kb) in control and HIRA KO livers. n = 2 per group (ATAC), n = 3 per group (CUT&Tag). E. Mean signal change (HIRA KO - control) for ATAC-seq, H3.3, H3K27ac, and H3K36me3 across all protein-coding genes (TSS to TES ± 3.0 Kb), grouped by expression deciles (D1, lowest; D10, highest). n = 2 per group (ATAC), n = 3 per group (CUT&Tag). F., G. Representative genome browser tracks at liver identity gene loci (Serpina6, Slc10a1) showing RNA-seq, ATAC-seq, H3.3, H3K27ac, and H3K36me3 signals in control and HIRA KO livers. Data are presented as means ± SEM. Statistical significance was determined by Pearson correlation (A) or unpaired Student’s t-test (B). Asterisk (*) indicates p < 0.05, (**) p < 0.01, (***) p< 0.001, (****) p < 0.0001.

To better understand the epigenetic mechanism underlying the loss of expression of HNF4α target genes and hepatocyte identity genes, we analyzed CUT&Tag of H3.3 (HIRA’s primary deposition substrate), H3K27ac (a mark of active enhancers and promoters), and H3K36me3 (a gene body-enriched mark deposited by SETD2 during transcription elongation that is critical to transcriptional fidelity and epigenetic stability^41,42^). As H3.3 differs from canonical H3.1/2 by only 4-5 amino acids, we first validated the H3.3 profiling approach^12^. Histone H3.3 is encoded by two genes, H3f3a and H3f3b, and combined conditional KO of both alleles has been previously generated^43^. Using liver tissue from an H3f3a^fl/fl^, H3f3b^fl/fl^ mouse injected with AAV8 Tbg-Cre to achieve hepatocyte-specific H3.3 deletion, we confirmed the expected reduction in H3.3 signal across all identified peaks (Suppl. Fig. 3F), validating the H3.3 CUT&Tag. Profile analysis from promoter to transcription end site (TES ± 3 kb) revealed that HIRA KO livers exhibit reduced H3.3 deposition across all protein-coding genes (Fig. 3C), as expected. H3K27ac and H3K36me3 were also both broadly reduced across all protein-coding genes in HIRA KO livers, including at liver identity genes (Fig. 3C, D). HIRA KO samples displayed reduced accessibility at the TSS of these identity genes (Fig. 3D), consistent with their tendency to decrease expression and loss of liver identity (Fig. 2C). Collectively, loss of identity and decreased expression of liver identity genes in HIRA KO associates with decreased TSS accessibility and decreased H3K27ac and H3K36me3 across genes.

Surprisingly, chromatin accessibility over gene bodies was modestly increased across all genes and markedly so at liver identity genes (Fig. 3C, D), despite their tendency toward decreased expression (Fig. 2C). This paradox suggests that HIRA KO disrupts gene expression and chromatin organization at identity genes in a non-canonical fashion. Because many liver identity genes are among the highly expressed genes (Suppl. Fig. 3G) and because genes downregulated by HIRA KO tend to be highly expressed (Fig. 1G), we examined whether the observed epigenetic changes were expression-dependent. We calculated the difference in signal between HIRA KO and control and grouped all genes by expression deciles. Disruption of chromatin accessibility and loss of H3.3, H3K27ac, and H3K36me3 signal, each scaled with baseline expression level, with the most highly expressed genes showing the greatest reduction (Fig. 3E). This indicates that HIRA-mediated chromatin maintenance is particularly critical at actively transcribed loci, consistent with elevated nucleosome turnover at highly expressed genes^44^. To illustrate these coordinated epigenetic changes at individual loci, we examined Serpina6 and Slc10a1, two representative liver identity genes (Fig. 3F, G). Each displayed reduced expression accompanied by decreased chromatin accessibility at promoters and putative enhancers, increased accessibility over gene bodies, and diminished H3.3, H3K27ac, and H3K36me3 signal, exemplifying the multi-layered epigenetic disruption caused by HIRA deletion. Taken together, these data demonstrate that HIRA maintains the hepatocyte epigenome linked to H3.3 deposition, with actively transcribed genes being most dependent. HIRA’s loss disrupts H3.3 deposition, chromatin accessibility, histone modifications, and DNA methylation, ultimately driving the erosion of liver cell identity and accelerated epigenetic aging.

### H3.3 knockout recapitulates HIRA-dependent gene expression changes but triggers acute hepatotoxicity

To test whether histone H3.3, rather than other H3.3-independent HIRA functions, mediates the effects of HIRA deletion on hepatocyte identity, we generated hepatocyte-specific H3.3 KO mice and performed parallel transcriptomic analyses. Following retro-orbital injection of AAV8 Tbg-Cre into 6-month-old H3f3a^fl/fl^, H3f3b^fl/fl^ animals, western blot analysis confirmed downregulation of H3.3 protein at three weeks post-injection (Fig. 4A). To assess the long-term consequences of hepatocyte H3.3 loss, we also aged H3.3 KO mice for one year (from 6 to18 months), mirroring our previous HIRA KO experiments. Surprisingly, in contrast to HIRA KO, aged animals showed complete restoration of H3.3 protein levels (Suppl. Fig. 4A), suggesting selective pressure against H3.3-deficient hepatocytes. Consistent with this interpretation, Cre recombinase protein was detectable in wildtype livers transduced with AAV Tbg-Cre but completely absent in aged H3.3 KO livers (Suppl. Fig. 4A), providing evidence that H3.3-deficient hepatocytes were eliminated. Body weight measurements at three weeks revealed no significant differences between H3.3 KO and control animals (Suppl. Fig. 4B). Likewise, in contrast to HIRA KO, plasma cholesterol and bile acid levels were not significantly altered in H3.3 KO mice (Suppl. Fig. 4C, D). However, assessment of liver damage through plasma enzyme levels demonstrated dramatic hepatotoxicity, with both ALT and ALP significantly elevated in H3.3 KO mice compared to controls (Fig. 4B, C). Together, these findings indicate that H3.3-deficient hepatocytes are likely markedly dysfunctional, incompatible with long-term hepatocyte survival, and replaced by compensatory proliferation of untransduced wildtype cells.

**Figure 4:**
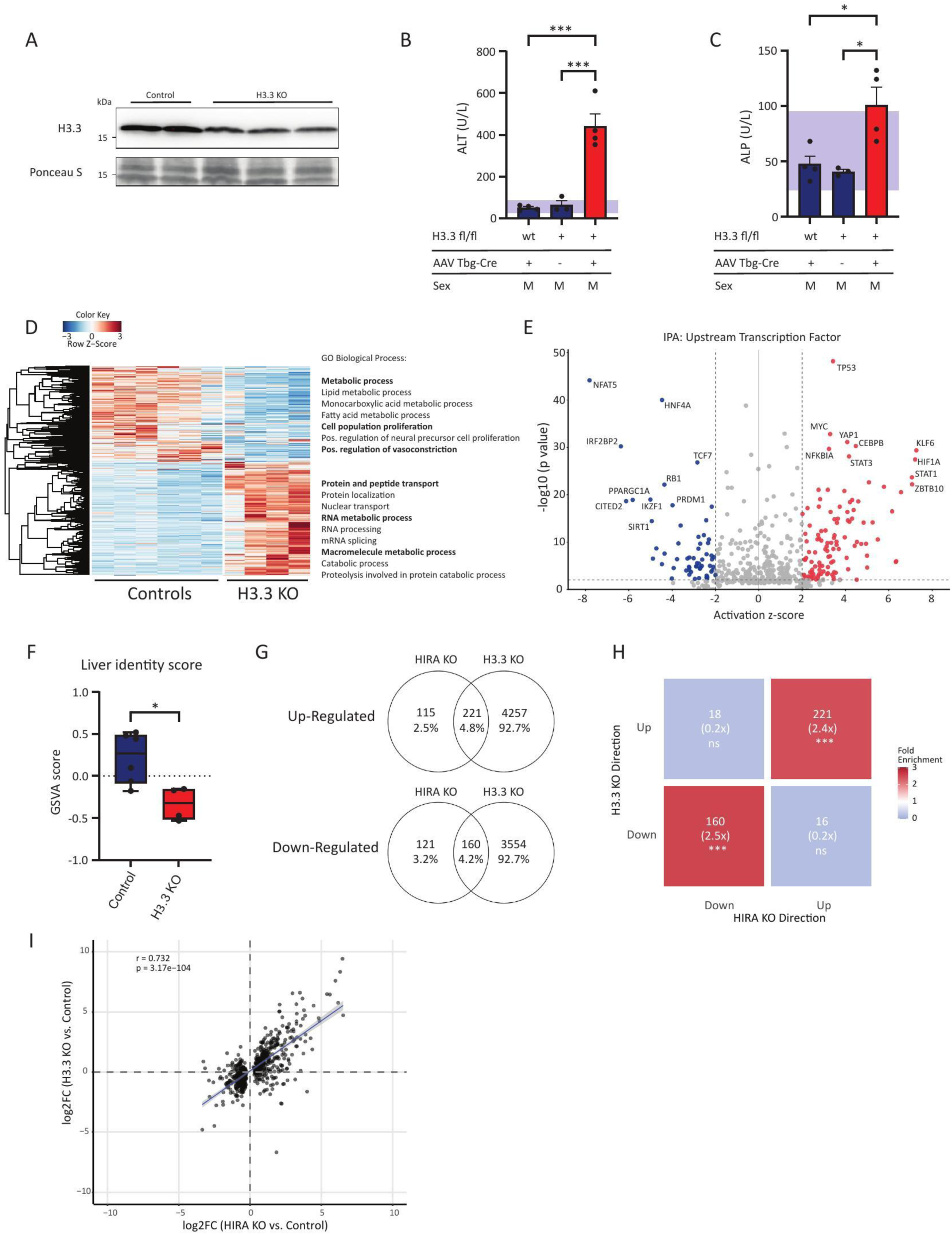
H3.3 knockout recapitulates HIRA-dependent gene expression changes but triggers acute hepatotoxicity. A. Representative western blot showing H3.3 protein levels in liver lysates from control and H3.3 KO male mice 3 weeks post-knockout. Ponceau S staining served as loading control. n = 3 (WT AAV Tbg-Cre, H3f3a^fl/fl^ H3f3b^fl/fl^ AAV ctl); n = 4 (H3f3a^fl/fl^ H3f3b^fl/fl^ AAV Tbg-Cre). B. Plasma alanine aminotransferase (ALT) and C. alkaline phosphatase (ALP) levels in control and H3.3 KO mice. D. Heatmap of DEGs from RNA-seq analysis comparing male controls with H3.3 KO mice. n = 3 (controls); n = 4 (HIRA KO). E. IPA showing top predicted upstream transcription factors regulating DEGs in control versus H3.3 KO mice. F. Gene set variation analysis (GSVA) of 85 liver identity genes in control and H3.3 KO mice. G. Venn diagrams showing the overlap of the DEGs in HIRA KO vs. H3.3 KO mice. H. Contingency analysis of DEGs shared between HIRA KO and H3.3 KO livers. Heatmap displays fold enrichment over expected overlap by chance for genes categorized by their direction of change in each knockout condition. Numbers indicate the count of overlapping genes, with fold enrichment shown in parentheses. I. Correlation between change in gene expression as a function of HIRA KO and change in gene expression as a function of H3.3 KO, among 523 DEGs with HIRA KO. n = 5 (HIRA KO); n = 3 (H3.3 controls); n = 4 (H3.3 KO). Data are presented as means ± SEM. Statistical significance was determined by one-way ANOVA with Tukey’s multiple comparisons test (B, C), unpaired Student’s t-test (F), Fisher’s exact test (H), or simple linear regression (I). Asterisk (*) indicates p < 0.05, (**) p < 0.01, (***) p< 0.001, (****) p < 0.0001.

In light of this eventual replacement of H3.3 KO hepatocytes, we performed transcriptomic analysis on male H3.3 KO livers three weeks post-injection, a timepoint when H3.3 KO was detectable (Fig. 4A), to compare the molecular consequences with HIRA deletion. PCA indicated that the control and H3.3 KO groups are distinctly separated (Suppl. Fig. 4E). RNA-sequencing identified 8,192 DEGs in H3.3 KO hepatocytes, substantially more than was observed in HIRA KO (617) (Fig. 4D). Gene Ontology analysis of upregulated transcripts revealed enrichment for ‘protein and peptide transport’, including ‘nuclear transport’, ‘RNA metabolic processes’ such as ‘mRNA splicing’, and pathways involved in ‘macromolecule metabolism’. In contrast, like HIRA KO, downregulated genes were predominantly associated with ‘metabolic processes’, particularly ‘lipid metabolic process’, ‘monocarboxylic acid metabolic process’, and ‘fatty acid metabolic process’ (Fig. 4D). IPA of upstream transcriptional regulators identified HNF4α as one of the most significantly inhibited factors, alongside other key metabolic transcription factors including PPARGC1A, and SIRT1 (Fig. 4E). Quantification of hepatocyte identity by GSVA analysis using the liver-specific gene signature demonstrated a significant reduction in the liver identity score in H3.3 KO animals (Fig. 4F), paralleling the phenotype observed in HIRA-deficient hepatocytes.

Direct comparison of the H3.3 KO and HIRA KO transcriptomes three weeks post KO revealed striking similarities. Venn diagram analysis showed 221 shared upregulated genes out of a total of 336 upregulated genes in HIRA KO (Fig. 4G). Out of the 281 downregulated transcripts in HIRA KO, 160 genes overlapped with H3.3 KO (Fig. 4G). Notably, this overlap was far greater than expected by chance. Directional overlap analysis showed that transcriptional changes induced by HIRA deletion and H3.3 loss were strongly concordant. Genes upregulated in both datasets (221; 2.4x over chance) and downregulated in both datasets (160; 2.5x over chance) were significantly enriched (Fig. 4H). In contrast, discordant gene regulation was rare and depleted: only 18 genes were down in H3.3 KO but up in HIRA KO, and 16 showed the opposite pattern (both 0.2x over chance, not significant; Fig. 4H). To further assess the directional concordance of expression changes, we plotted the log2 fold changes of all DEGs from the 3-week HIRA KO dataset against their corresponding expression changes in 3-week H3.3 KO livers. This analysis revealed a strong positive correlation (r = 0.732, Fig. 4I). In sum, although H3.3 KO triggered more severe phenotypic disruption than HIRA KO, the transcriptional changes at 3 weeks strongly phenocopied those of HIRA KO, indicating that the consequences of HIRA KO are likely mediated by a block to H3.3 deposition.

### Liver regeneration rescues the phenotypic consequences of HIRA loss through proliferation-dependent mechanisms

The data so far demonstrate that HIRA KO disrupts epigenetic maintenance and cell identity in non-proliferating hepatocytes. However, replication-coupled histone deposition during cell division could provide an alternative mechanism to restore chromatin integrity. Therefore, we set out to ask whether defects in epigenetic maintenance and cell identity caused by HIRA KO could be rescued by reactivation of cell proliferation. To do this, we induced acute hepatocyte proliferation by triggering liver regeneration via partial hepatectomy after KO of HIRA in adult mice. Mouse livers can nearly fully regenerate within 5-7 days, with Ki67 expression peaking around days 2-3 post-surgery^45^. Using the two already introduced approaches (Cre-mediated in HIRA^fl/fl^ and CRISPR-mediated in wildtype mice), we deleted HIRA in hepatocytes of 6-month-old mice, waited four weeks, induced regeneration by partial hepatectomy, and collected livers five days later (Fig. 5A). Indicative of effective regeneration, liver-to-body weight ratios had recovered five days after hepatectomy (Fig. 5B). Measuring the regenerating lobes specifically (right and caudate) showed the expected increase in lobe-to-body weight ratio after hepatectomy, confirming robust compensatory growth in both genotypes (Fig. 5C). Representative lobe images with and without regeneration are shown in Supplementary Fig. 5A. In these analyses, we detected no significant difference between extent of regeneration of control and HIRA KO livers, detected either by weight or appearance. We also confirmed by Western blot that HIRA levels were diminished in HIRA KO before and after liver regeneration, and that H3.3 was reduced in HIRA KO livers before regeneration (Fig. 5D, Suppl. Fig. 5B). Strikingly, after regeneration, H3.3 was strongly diminished in both control and KO livers, while total H3 remained stable (Fig. 5D). This was reflected in an increased ratio of total H3/ H3.3 in both control and HIRA KO after regeneration, suggesting replacement of H3.3 by canonical H3.1/2 (Fig. 5D, Suppl. Fig. 5B).

**Figure 5:**
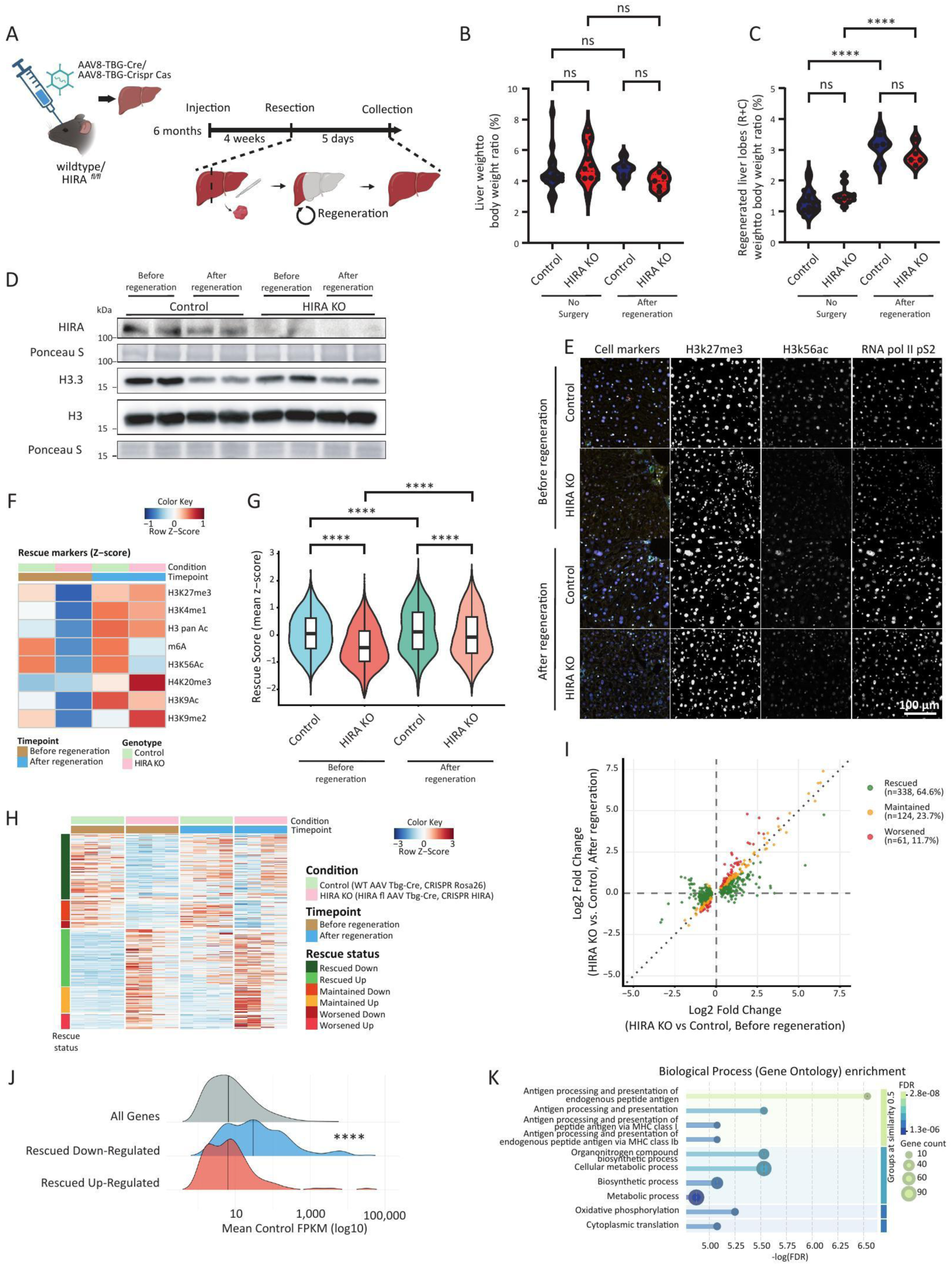
Liver regeneration rescues the phenotypic consequences of HIRA loss through proliferation-dependent mechanisms. A. Graphical representation of the experimental design for hepatectomy-induced liver regeneration. B. Liver weight to body weight ratio of control and HIRA KO liver without surgery and after regeneration. n = 17 (control without surgery); n = 12 (HIRA KO without surgery); n = 8 (control after regeneration); n = 5 (HIRA KO after regeneration). C. Regenerated liver lobes (right (R) and caudate (C) lobes) weight to body weight ratio without surgery and after regeneration. D. Representative western blot showing HIRA, H3.3 and total H3 protein levels in liver lysates from control and HIRA KO male mice before and after regeneration. Ponceau S staining served as loading control. E. Representative CODEX multiplex immunofluorescence images of liver tissue from control and HIRA KO mice before and after partial hepatectomy-induced regeneration. Lineage markers (HNF4α (red), hepatocytes; CK19 (green), cholangiocytes; CD45 (yellow), immune cells) are shown with DAPI (blue; nuclei) and β-catenin (white; membrane). Grayscale panels show H3K27me3, H3K56ac, and RNA Pol II pS2. Scale bar, 100 μm. F. Z-scored heatmap of rescue markers in hepatocytes (HNF4α^+^, CK19^−^, CD45^−^; n = 194 165 cells) across conditions and timepoints. Rescue markers represent epigenetic modifications that were altered by HIRA loss and shifted toward control levels after regeneration. G. Quantification of composite rescue score (mean z-score of rescue markers) in hepatocytes from control and HIRA KO livers before and after regeneration. H. Heatmap of RNA-seq data comparing male control (WT AAV Tbg-Cre, WT AAV CRISPR Rosa26) with HIRA KO (HIRA^fl/fl^ AAV Tbg-Cre, WT AAV CRISPR HIRA) mice before and after regeneration. The DEGs comparing control and HIRA KO samples before regeneration are presented and subsequently clustered based on their rescue status after regeneration. Genes exhibiting more than 20% upregulation (control versus HIRA KO after regeneration) following initial downregulation, or genes showing more than 20% downregulation after initial upregulation, were classified as rescued. n = 4 per group (2 mice per knockout approach). I. Correlation between change in gene expression as a function of HIRA KO before regeneration and change in gene expression as a function of HIRA KO after regeneration, among 523 DEGs with HIRA KO. J. Ridgeline plot illustrating expression distribution of all expressed genes compared to the rescued significantly up- and down-regulated DEGs. K. Gene Ontology (GO) Biological Process enrichment analysis of downregulated rescued genes. Top 10 enriched terms are shown, sorted by −log(FDR) and grouped by semantic similarity (≥0.5). Dot size represents gene count; color indicates FDR. Data are presented as means ± SEM. Statistical significance was determined by two-way ANOVA with Tukey’s multiple comparisons test (B, C, G), or Wilcoxon rank-sum test (Mann-Whitney U test) (J). Asterisk (*) indicates p < 0.05, (**) p < 0.01, (***) p< 0.001, (****) p < 0.0001.

To extend this epigenetic analysis, we assessed whether regeneration restores epigenetic signatures at single-cell and spatial resolution using multiplex CODEX imaging with a 35-marker epigenetic panel supplemented with lineage markers (HNF4α for hepatocytes, CK19 for cholangiocytes, and CD45 for immune cells) (Fig. 5E-G; Suppl. Fig. 5C-F). Representative images show the cell-type markers together with selected chromatin and transcription marks (H3K27me3, H3K56ac, and RNA Pol II pS2) across genotypes and timepoints (Fig. 5E). Unsupervised clustering using the full marker set (cell identity, chromatin and transcription) resolved distinct cellular populations, visualized by UMAP embedding and marker dot plots (Suppl. Fig. 5C, D). Restricting analysis to hepatocytes (HNF4α+, CK19-, CD45-), we separated epigenetic features into (i) marks that were altered by HIRA loss before regeneration and shifted toward control levels after regeneration (“rescue markers”), (ii) marks that primarily changed with regeneration in both genotypes (“regeneration markers”), and (iii) marks that were largely stable (“no change”) (Suppl. Fig. 5E). The rescue markers are shown as a z-scored heatmap (Fig. 5F) and quantified as a composite rescue score, which showed a significant restoration in HIRA KO hepatocytes after regeneration (Fig. 5G). Finally, mapping the rescue score back into tissue space did not reveal an obvious zonation pattern, suggesting that the regeneration-associated epigenetic normalization we observe is broadly distributed (Suppl. Fig. 5F).

To determine whether replacement of H3.3 by H3.1/2 and the restoration of chromatin signatures captured by multiplex CODEX imaging were reflected at the transcriptional level, we performed RNA-seq from control and HIRA KO livers before and after regeneration. Samples separated appropriately by condition and timepoint using PCA (Suppl. Fig. 5G). Proliferation after hepatectomy was independently confirmed by enrichment of mitotic cell cycle gene sets in both control and HIRA KO livers (Suppl. Fig. 5H, I). We then focused on the 523 DEGs identified between control and HIRA KO livers prior to regeneration and tracked their expression after regeneration. These DEGs were clustered across conditions and classified according to their expression pattern: ‘Up’ or ‘Down’, according to increased or decreased expression in HIRA KO compared to control; ‘Rescued’, ‘Maintained’ or ‘Worsened’, according to the effect of regeneration on the difference between HIRA KO and control (Fig. 5H). For genes up in HIRA KO, many differences were reduced by regeneration (‘Rescued Up’), although this was largely due to increased expression in regenerating controls (Fig. 5H). However, consistent with the epigenetic analyses, regeneration broadly restored expression of many genes down in HIRA KO (‘Rescued Down’, Fig. 5H). Plotting the fold change of HIRA KO DEGs pre-regeneration against their corresponding fold change post-regeneration demonstrated that most DEGs showed reduced differences following regeneration, with the majority classified as rescued (Fig. 5I). Notably, this analysis also highlighted that a subset of genes remained persistently altered or diverged further (maintained and worsened) after regeneration, emphasizing that rescue is substantial but not uniform across all HIRA-sensitive transcripts (Fig. 5I). To contextualize which classes of genes were most affected, we compared expression distributions and found that rescued downregulated genes were significantly enriched for highly expressed liver genes compared to all detected genes and upregulated genes (Fig. 5J). Gene ontology analysis of the rescued downregulated cluster showed enrichment for core liver functional programs such as metabolic and biosynthetic processes, alongside immune-related terms including antigen processing and presentation of endogenous antigen (Fig. 5K).

Our data demonstrate that liver regeneration mitigates many of the epigenetic and transcriptional defects in HIRA KO livers, suggesting that replication-coupled histone incorporation may retroactively compensate, at least in part, for the absence of HIRA-mediated chromatin assembly.

## Discussion

Epigenetic drift has been proposed to drive tissue dysfunction during aging^46–48^. However, this model likely applies differently to proliferating and non-proliferating cells. A fundamental challenge in aging tissues is how postmitotic cells preserve chromatin integrity and cellular identity over time^49,50^. Even in non-proliferating cells, chromatin is dynamic and plastic, continually disassembled and re-assembled in association with transcription and DNA repair. While proliferating cells renew chromatin states during DNA replication largely through CAF-1-mediated deposition of canonical H3.1 and H3.2 histones, distribution of parental histones between daughter strands, and copying of histone and DNA modifications, postmitotic hepatocytes must maintain their chromatin landscape without DNA synthesis^51–53^. Our data show that HIRA-mediated, replication-independent H3.3 deposition fulfills this role, predominantly at highly expressed genes that require frequent nucleosome disassembly and reassembly during active transcription. Loss of HIRA leads to progressive chromatin deterioration, including changes in accessibility, DNA methylation, histone modifications, and ultimately results in transdifferentiation toward a cholangiocyte-like state. HIRA deletion accelerates epigenetic age and promotes fibrosis in an age-dependent manner, with young HIRA KO showing transcriptional changes but minimal pathology, whereas aged KOs develop severe fibrosis, suggesting that chromatin defects sensitize tissues to age-related insults.

Mechanistically, the preferential downregulation of highly expressed genes in HIRA KO hepatocytes suggests that these loci are particularly dependent on replication-independent H3.3 deposition to maintain chromatin integrity. Without HIRA, these loci are vulnerable to progressive chromatin deterioration, with consequences across multiple epigenetic layers, including altered histone modifications, aberrant DNA methylation, and changes in chromatin accessibility. In contrast, upregulated genes in HIRA KO livers likely represent secondary consequences of identity loss rather than direct targets of HIRA-mediated maintenance, consistent with the temporal dissociation between early transcriptional changes and the later emergence of fibrosis and transdifferentiation in aged knockouts. Collectively, this supports a model in which failed chromatin maintenance at identity genes leads to broad epigenetic erosion and ultimately lineage infidelity, underscoring that chromatin homeostasis is not a passive state but an active, ongoing requirement in terminally differentiated cells.

The tight coupling between hepatocyte identity loss and metabolic dysfunction in our model highlights the inseparable nature of cell-type specification and metabolic specialization in hepatocytes. This connection arises for multiple reasons. First, as the major metabolic organ, the liver’s identity is largely defined by metabolic genes, and since HIRA preferentially maintains highly expressed hepatocyte identity genes, many of these are metabolic in nature. Second, chromatin state and metabolism form interdependent networks; chromatin structure influences metabolic gene expression, while metabolic intermediates, such as acetyl-CoA, serve as substrates for chromatin-modifying enzymes^54,55^. Thus, disruption of chromatin maintenance through HIRA loss simultaneously erodes both cellular identity and metabolic function.

The partial rescue of HIRA KO epigenome and transcriptome through liver regeneration demonstrates that chromatin states in differentiated cells retain plasticity and are not irreversibly determined. During the regeneration process, replication-coupled histone deposition appears capable of re-establishing much of the normal hepatocyte chromatin landscape, consistent with recent work by other groups^56–58^. This implies that cells retain sufficient epigenetic information to restore chromatin states when provided with appropriate cues such as DNA replication, supporting the concept of a retrievable “back-up copy” of epigenetic identity^7^. Analysis of epigenetic markers following regeneration revealed rescue of both activating and repressive histone modifications, suggesting that replication-coupled mechanisms can broadly restore the chromatin landscape. However, some transcriptional defects persist even after regeneration. The imperfect rescue likely reflects heterogeneity in the proliferative response, where some cells divide more extensively than others, and others may exit the cell cycle early after initial division. Time-course analysis of regeneration dynamics could provide further insight into how chromatin states are progressively restored and whether iterative rounds of replication are required for full epigenetic normalization.

The phenomenon of cell identity loss has recently been recognized as a common feature of various diseases and aging: Studies in cancer have shown that loss of lineage-specific transcription factor expression leads to dedifferentiation and metastatic potential^59,60^. Studies in diabetes have demonstrated that pancreatic beta cells lose their identity markers under metabolic stress^61,62^. Studies in neurodegenerative diseases show progressive loss of cell-type-specific gene expression programs^63^. Our findings suggest that defects in chromatin maintenance machinery can be a common upstream mechanism driving identity loss, and thus raise the questions of whether identity loss serves as a precursor to disease and whether reinforcing cell identity is a potential early intervention point before onset of irreversible pathology. If chromatin maintenance defects contribute to age-related tissue dysfunction, therapeutic approaches that protect cell identity warrant exploration. HNF4α, which is downregulated in both acute and chronic liver injury, represents a therapeutic target for enhancing liver function through identity restoration^64^. Indeed, proof-of-concept studies in liver disease have demonstrated that restoring the master hepatocyte transcription factor HNF4α via mRNA delivery in lipid nanoparticles can reverse fibrosis in preclinical models by re-establishing hepatocyte metabolic function and identity^65^. Beyond liver-directed approaches, transcription factor-based interventions building on pioneering reprogramming work offer additional avenues^66^. Recent cellular reprogramming studies show progressive epigenetic resetting, yet the mechanisms ensuring correct tissue-specific chromatin landscape re-establishment remain incompletely understood^67–69^. Our regeneration experiments suggest that cell division may help reset aberrant chromatin states. Understanding how cells navigate between plasticity during reprogramming and stability during differentiation will be crucial for developing safe therapeutic approaches.

A critical question emerging from this work concerns why aged livers still exhibit epigenetic erosion and identity loss despite HIRA’s presence^45,70^. We hypothesize that age-related epigenetic damage may accumulate faster than HIRA can sustain chromatin stability. This is similar to how DNA damage accumulates with age when repair mechanisms can no longer keep pace with the rate of damage accrual^71^. It highlights a fundamental principle in aging biology: protective mechanisms that function effectively in youth become overwhelmed during aging when the rate of damage exceeds their maintenance capacity. However, several key mechanistic questions remain unresolved. Whether H3.3 functions primarily as a substrate for replication-independent deposition or plays a more instructive role through variant-specific properties remains an active area of investigation. Recent work has identified H3.3 Ser31 phosphorylation as a unique regulatory mechanism that modulates nucleosome dynamics and transcriptional responses, suggesting functional roles beyond deposition alone^72–74^. Notably, the more severe phenotype of H3.3 KO compared to HIRA KO implies that H3.3 has additional HIRA-independent functions, likely mediated in part through DAXX/ATRX-dependent deposition at heterochromatic regions and telomeres^11^. Another future goal is to determine if metabolic dysfunction and fibrosis can be separated from identity loss or represent inevitable consequences of hepatocyte dedifferentiation. Validating these findings beyond the liver and testing these phenomena in human tissues will be crucial, despite the challenges in obtaining suitable samples.

In conclusion, our work establishes that differentiated cell identity requires active chromatin maintenance during aging through HIRA-mediated H3.3 deposition. Loss of this maintenance allows chromatin dysregulation, metabolic dysfunction, and liver pathology. These findings reframe our understanding of how cell identity is preserved in adult tissues and show that active chromatin homeostasis over the lifespan is necessary to prevent tissue dysfunction. The partial reversibility of these defects through cell division offers an approach for regenerative interventions while highlighting the critical importance of dedicated chromatin maintenance mechanisms in postmitotic cells. As we develop therapeutic approaches for aging and regenerative medicine, maintaining proper cell identity through chromatin homeostasis may prove to be as important as replacing lost cells.

## Methods

**Suppl. Table 1:** CosMX fibrosis score markers.

| Gene No. | Marker |
| --- | --- |
| 1 | Col1a1 |
| 2 | Col1a2 |
| 3 | Col3a1 |
| 4 | Col4a1 |
| 5 | Col5a1 |
| 6 | Col6a1 |
| 7 | Fn1 |
| 8 | Vcan |
| 9 | Mmp14 |
| 10 | Mmp2 |
| 11 | Mmp7 |
| 12 | Mmp9 |
| 13 | Timp1 |
| 14 | Col1a1 |
| 15 | Col3a1 |
| 16 | Fn1 |
| 17 | Lcn2 |
| 18 | Tgfbr2 |
| 19 | Col1a1 |
| 20 | Col3a1 |
| 21 | Mmp2 |
| 22 | Pdgfrb |
| 23 | Timp1 |
| 24 | Cxcl14 |
| 25 | Itgb6 |
| 26 | Muc1 |
| 27 | Ccl2 |
| 28 | Cd68 |
| 29 | Il1b |
| 30 | Lgals3 |
| 31 | Spp1 |
| 32 | Tnf |
| 33 | Acta2 |
| 34 | Fap |
| 35 | Myh11 |
| 36 | Pdgfra |
| 37 | Pdgfrb |
| 38 | Tagln |
| 39 | Smad2 |
| 40 | Smad3 |
| 41 | Tgfb1 |
| 42 | Tgfbr1 |
| 43 | Tgfbr2 |

### Antibodies

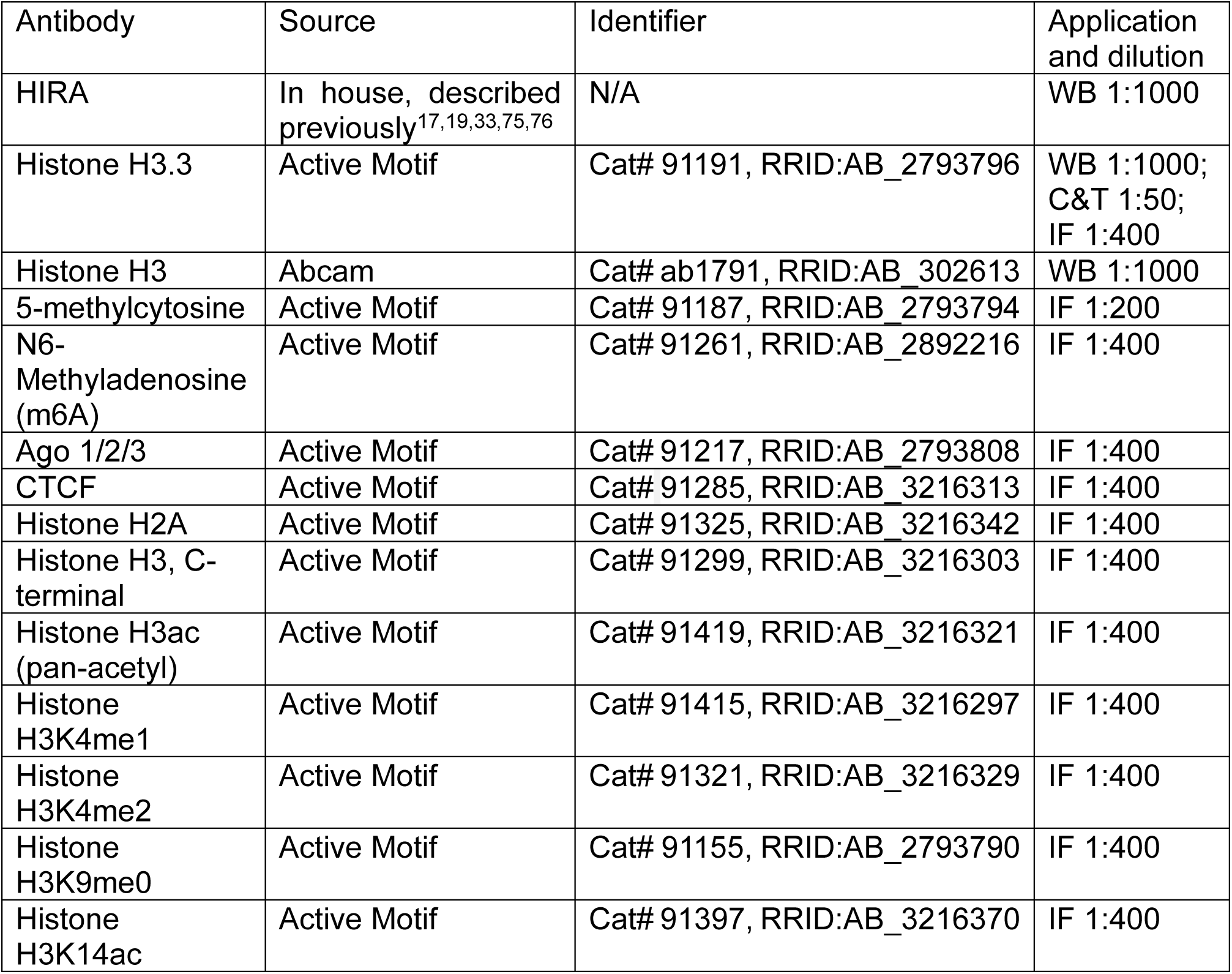
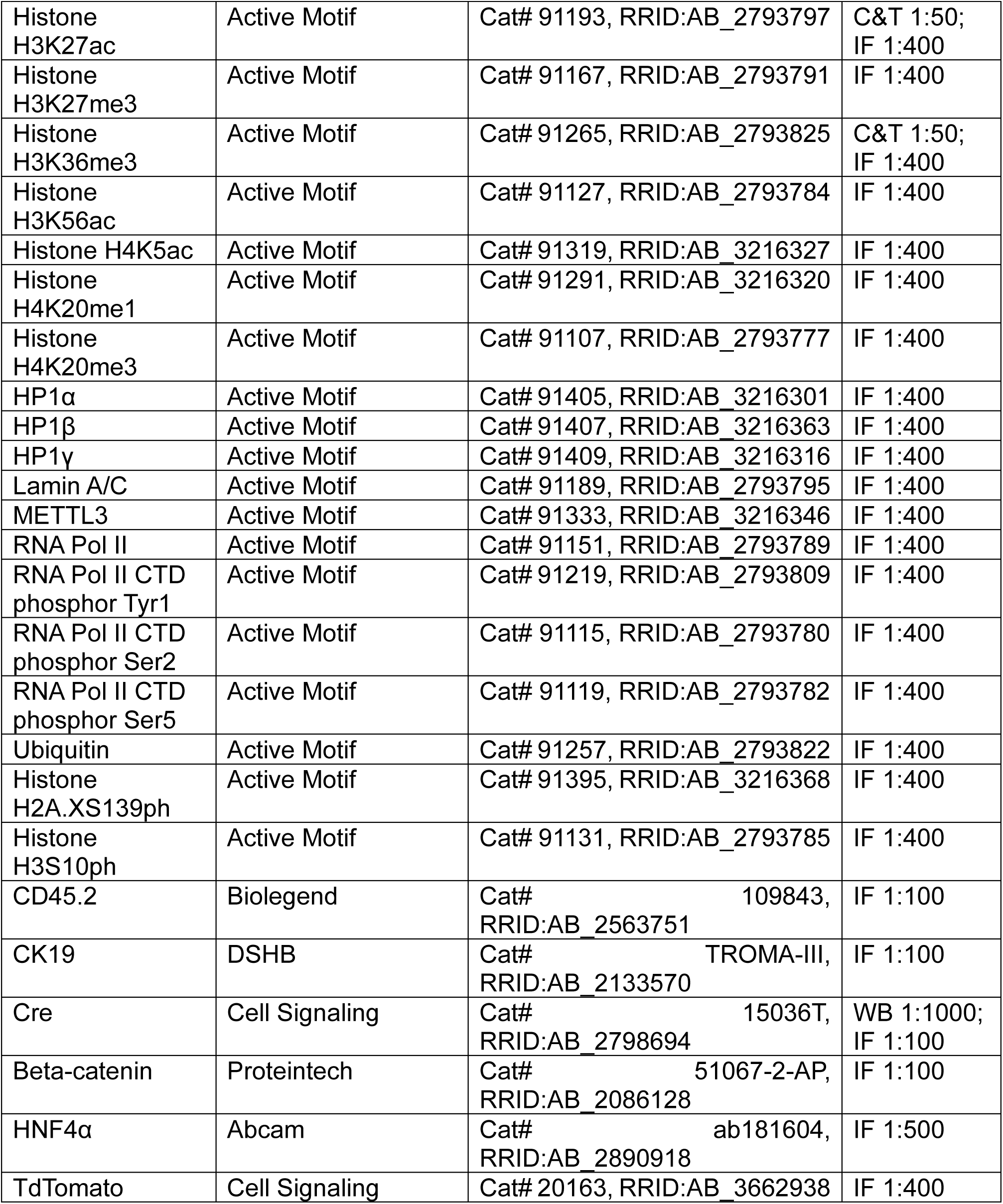

| Antibody | Source | Identifier | Application and dilution |
| --- | --- | --- | --- |
| HIRA | In house, described previously<sup>17,19,33,75,76</sup> | N/A | WB 1:1000 |
| Histone H3.3 | Active Motif | Cat# 91191, RRID:AB_2793796 | WB 1:1000;<br>C&T 1:50;<br>IF 1:400 |
| Histone H3 | Abcam | Cat# ab1791, RRID:AB_302613 | WB 1:1000 |
| 5-methylcytosine | Active Motif | Cat# 91187, RRID:AB_2793794 | IF 1:200 |
| N6-Methyladenosine (m6A) | Active Motif | Cat# 91261, RRID:AB_2892216 | IF 1:400 |
| Ago 1/2/3 | Active Motif | Cat# 91217, RRID:AB_2793808 | IF 1:400 |
| CTCF | Active Motif | Cat# 91285, RRID:AB_3216313 | IF 1:400 |
| Histone H2A | Active Motif | Cat# 91325, RRID:AB_3216342 | IF 1:400 |
| Histone H3, C-terminal | Active Motif | Cat# 91299, RRID:AB_3216303 | IF 1:400 |
| Histone H3ac (pan-acetyl) | Active Motif | Cat# 91419, RRID:AB_3216321 | IF 1:400 |
| Histone H3K4me1 | Active Motif | Cat# 91415, RRID:AB_3216297 | IF 1:400 |
| Histone H3K4me2 | Active Motif | Cat# 91321, RRID:AB_3216329 | IF 1:400 |
| Histone H3K9me0 | Active Motif | Cat# 91155, RRID:AB_2793790 | IF 1:400 |
| Histone H3K14ac | Active Motif | Cat# 91397, RRID:AB_3216370 | IF 1:400 |
| Histone H3K27ac | Active Motif | Cat# 91193, RRID:AB_2793797 | C&T 1:50;<br>IF 1:400 |
| Histone H3K27me3 | Active Motif | Cat# 91167, RRID:AB_2793791 | IF 1:400 |
| Histone H3K36me3 | Active Motif | Cat# 91265, RRID:AB_2793825 | C&T 1:50;<br>IF 1:400 |
| Histone H3K56ac | Active Motif | Cat# 91127, RRID:AB_2793784 | IF 1:400 |
| Histone H4K5ac | Active Motif | Cat# 91319, RRID:AB_3216327 | IF 1:400 |
| Histone H4K20me1 | Active Motif | Cat# 91291, RRID:AB_3216320 | IF 1:400 |
| Histone H4K20me3 | Active Motif | Cat# 91107, RRID:AB_2793777 | IF 1:400 |
| HP1α | Active Motif | Cat# 91405, RRID:AB_3216301 | IF 1:400 |
| HP1β | Active Motif | Cat# 91407, RRID:AB_3216363 | IF 1:400 |
| HP1γ | Active Motif | Cat# 91409, RRID:AB_3216316 | IF 1:400 |
| Lamin A/C | Active Motif | Cat# 91189, RRID:AB_2793795 | IF 1:400 |
| METTL3 | Active Motif | Cat# 91333, RRID:AB_3216346 | IF 1:400 |
| RNA Pol II | Active Motif | Cat# 91151, RRID:AB_2793789 | IF 1:400 |
| RNA Pol II CTD phosphor Tyr1 | Active Motif | Cat# 91219, RRID:AB_2793809 | IF 1:400 |
| RNA Pol II CTD phosphor Ser2 | Active Motif | Cat# 91115, RRID:AB_2793780 | IF 1:400 |
| RNA Pol II CTD phosphor Ser5 | Active Motif | Cat# 91119, RRID:AB_2793782 | IF 1:400 |
| Ubiquitin | Active Motif | Cat# 91257, RRID:AB_2793822 | IF 1:400 |
| Histone H2A.XS139ph | Active Motif | Cat# 91395, RRID:AB_3216368 | IF 1:400 |
| Histone H3S10ph | Active Motif | Cat# 91131, RRID:AB_2793785 | IF 1:400 |
| CD45.2 | Biolegend | Cat# 109843, RRID:AB_2563751 | IF 1:100 |
| CK19 | DSHB | Cat# TROMA-III, RRID:AB_2133570 | IF 1:100 |
| Cre | Cell Signaling | Cat# 15036T, RRID:AB_2798694 | WB 1:1000;<br>IF 1:100 |
| Beta-catenin | Proteintech | Cat# 51067-2-AP, RRID:AB_2086128 | IF 1:100 |
| HNF4α | Abcam | Cat# ab181604, RRID:AB_2890918 | IF 1:500 |
| TdTomato | Cell Signaling | Cat# 20163, RRID:AB_3662938 | IF 1:400 |

### Plasmids

|  |  |
| --- | --- |
| 1 | AAV-TBG-MCS_108bp (produced by Vector Builder) |
| 2 | AAV-TBG-PI-Cre (Addgene #107787) |
| 3 | pX602-AAV-TBG-SaCas9-U6-sgRosa26 |
| 4 | pX602-AAV-TBG-SaCas9-U6-sgHIRA |

### Cas9 guide RNA sequences

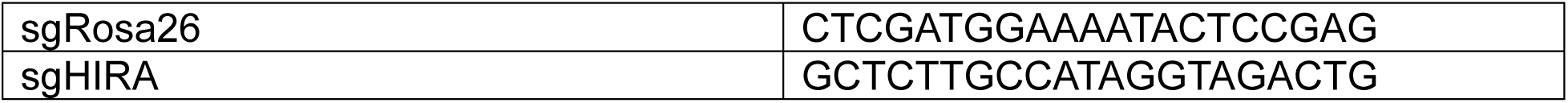

### Mice

All animal procedures were approved by the Institutional Animal Care and Use Committee (IACUC) of Sanford Burnham Prebys Medical Discovery Institute. Animal experiments were performed at Sanford Burnham Prebys Medical Discovery Institute Animal Facility in compliance with the IACUC guidelines. The studies performed within this manuscript were performed with all relevant ethical regulations regarding animal research. Young and old C57BL/6J animals were obtained from the NIA aging colony housed at Charles Rivers. HIRA^fl/fl^, H3f3a^fl/fl^ H3f3b^fl/fl^, LSL-tdTomato mice were all previously generated and described^33,43,77^. Mice were genotyped both by PCR analysis and Transnetyx (www.transnetyx.com). Animals were housed 5 mice per cage and maintained under controlled temperature (22.5°C) and illumination (12h dark/light cycle) conditions. Teklad Global 18% Protein Rodent diet was provided ad libitum.

### Western blot

Snap-frozen liver tissues were lysed using a bead tube homogenizer in SDS sample buffer (62.5 mM Tris at pH 6.8, 0.01% bromophenol blue, 10% glycerol, and 2% SDS) supplemented with protease and phosphatase inhibitor. To further reduce genomic DNA contamination and other debris, samples were sonicated and subsequently centrifuged (15,000 x g, 10 min, 4°C) to pellet insoluble material. Next, lysates were incubated at 95°C for 5 min. The proteins were then separated on precast protein gels (4-20% Mini-PROTEAN^®^ TGX™) and transferred onto nitrocellulose membranes. Ponceau S stain was taken. To detect the proteins of interest, membranes were blocked for 1 h in 5% non-fat milk supplemented with 0.1% Tween 20 (TBST) and were then incubated overnight at 4 °C with primary antibodies. Primary antibodies were diluted in blocking solution. After 3x washing in TBS Tween, the membranes were incubated for 1 h with the corresponding secondary antibody conjugated with horseradish peroxidase. After another 3x washing steps with TBS Tween, the signal was detected using either SuperSignal™ West Pico or Femto™ Substrate (Thermo Scientific) and a BioRad ChemiDoc™. Densitometry was performed using ImageJ software. Blots were quantified by normalizing to Ponceau S. All information regarding the antibodies used is provided in the supplemental information.

### Immunofluorescence

Liver tissues were fixated for 24 h at room temperature with 10% Neutral Buffered Formalin. Following fixation, the tissues were paraffin embedded and sectioned into 5 µm sections. Sections were deparaffinized with 2x 5 min washes in xylene and rehydrated through graded ethanol to water (3 min 95% Ethanol, 3 min 90% Ethanol, 3 min 70% Ethanol, 2 min 50% Ethanol). Antigen retrieval was performed in citrate buffer pH 6.0 using a steamer for 30 min, followed by cooling to room temperature and 15 min permeabilization in 0.05% Triton X-100 PBS. Next, the tissues were blocked in 4% BSA, 1% goat serum and 1% FBS at room temperature for 2 h. After blocking, the sections were incubated with primary antibodies in blocking buffer overnight at 4°C, washed three times with PBS + 1% Triton X-100 and then incubated with secondary antibodies in blocking buffer for 1 h at room temperature. After another three washes with PBS + 1% Triton X-100, Vector TrueVIEW® Autofluorescence Quenching Kit was used according to manufacturer’s protocol. DAPI was applied at 5 µg/ml for 10 min and subsequently washed 3x in PBS before coverslips were mounted in Fluoromount-G™. Images were taken using a Nikon Eclipse Ti2 epifluorescence microscope and data were processed with Nikon NIS-Elements software.

### Liver regeneration

70% partial hepatectomy (PHx) was conducted according to the protocol described by Mitchell and Willenbring^78^. Briefly, the left lateral and median lobes were surgically resected and collected as baseline (“before regeneration”) tissue. Mice were allowed to recover for five days post-surgery, after which they were euthanized via carbon dioxide inhalation followed by cervical dislocation. Regenerated liver tissue was harvested and either snap-frozen in liquid nitrogen, OCT (Tissue-Tek) embedded, or fixed and processed for paraffin embedding. To minimize variability associated with circadian influences on hepatocyte proliferation, all surgical procedures were performed during the morning hours.

### SaCas9 sgRNA design and validation

Single guide RNAs (sgRNAs) targeting genes of interest were designed using the CRISPick algorithm against the mouse reference genome GRCm38 (Ensembl v.102). Selected sgRNA sequences were cloned into the pX602-AAV-TBG::NLS-SaCas9-NLS-HA-OLLAS-bGHpA;U6::BsaI-sgRNA vector and transformed into NEB Stable competent *E. coli*. Successful sgRNA insertion was confirmed by Sanger sequencing of bacterial colonies (Genewiz) using a universal U6 promoter primer (5’-GACTATCATATGCTTACCGT-3’). To verify editing efficiency *in-vivo*, genomic DNA was isolated from mouse liver tissue using the Monarch Genomic DNA Purification Kit. A ∼500 bp region flanking the predicted sgRNA cut site was amplified by PCR with 100 ng of template gDNA and 1 µL of 100 µM forward and reverse primers. Amplicons were purified with the QIAquick PCR Purification Kit and then sequenced (Genewiz). Indel frequencies and knockout efficiency were quantified using Inference of CRISPR Edits (ICE) analysis (Synthego ICE v3.0).

### AAV production

Recombinant adeno-associated virus (rAAV) was produced at large scale by the SBP Functional Genomics Core Facility. HEK-293T cells were co-transfected with the rAAV vector, RepCap8, and pHelper plasmids using polyethylenimine. 96 h post-transfection, viral particles were collected from the culture supernatant by polyethylene glycol precipitation and from cell pellets by sonication, followed by benzonase digestion. Viral particles were then purified via iodixanol density gradient ultracentrifugation. The rAAV-containing fraction was collected, buffer-exchanged into PBS, aliquoted, and stored at −80 °C. Viral titers were quantified by SYBR Green qPCR. Virus suspension was injected at a dose of 3.33 × 10^11^ viral genomes per mouse in 100 µl sterile PBS retro-orbitally.

### CosMx

The median lobe was harvested, fixed in 10% Neutral Buffered Formalin and processed for paraffin embedding. The 1000-plex Mouse Universal Cell Characterization RNA Panel was applied to 5 µm sections, and spatial profiling was conducted by Bruker Spatial Biology (Seattle, WA). Files were processed using Seurat (v4.9.9.9050). Three biological replicates were analyzed, with 7-9 fields of view (FOVs) per tissue. All cells within these FOVs were combined to create a composite for analysis. Cells falling within the top or bottom 15^th^ percentile for detected features, or with total counts below 20, were excluded from downstream analysis. Filtered samples were normalized using SCTransform (v0.3.5) with a clip range of −10 to 10, and batch-integrated using Harmony (v1.2.3). Cell type annotations were assigned based on the top 5 genes identified through unbiased cluster-specific gene expression analysis, ranked by log2 fold change. Annotations were validated using established liver cell type markers: hepatocytes (*Apoa1*, *Glul*, *Serpina1a*, *Apoe*), cholangiocytes (*Spp1*, *Krt19*, *Epcam*, *Sox9*), hepatic stellate cells (*Dcn*, *Bmp5*, *Col1a1*, *Col1a2*), Kupffer cells (*Cd5l*, *Cd74*, *C1qa*, *C1qb*), endothelial cells (*Lyve1*), T cells (*Cd2*, *Cd3d*, *Cd3g*), NK cells (*Stat4*, *Nkg7*), B cells (*Cd79a*), and pan-immune cells (*Ptprc*). Neighborhood was defined as the 20 nearest neighbors of cholangiocytes.

### Multiplex Immunofluorescence (CODEX)

Liver tissues collected before and after regeneration were embedded in OCT and rapidly frozen by immersion in liquid nitrogen-cooled isopentane. Tissue blocks were stored at −80 °C until sectioning. Cryosections (8 μm) were prepared, stained, and imaged using the Akoya Biosciences Phenocycler system as previously described^79^. Epigenetic antibodies (Active Motif AbFlex) and cell type markers were conjugated to unique DNA barcodes for multiplexed detection with copper-free click chemistry by DBCO addition to the antibody and SPAAC reaction with an azide-conjugated DNA oligonucleotide^80^. DBCO introduction was achieved via Sortase enzymatic reaction of the AbFlex antibodies (bearing Sortase recognition sequence) with poly Gly probe comprising three DBCO moieties. This approach allows site specific (to Fc region of AbFlex) conjugation of multiple SPAAC counterparts followed by reaction with azide modified oligonucleotides (Sortag-IT 3X DBCO Click Labeling kit).

Multiplex images were processed using CRISP (Convolutional Registration of Images with Subpixel Precision) for tile alignment, drift compensation, deconvolution, and stitching^79^. Background autofluorescence was subtracted using blank cycles imaged at the beginning and end of each run. Cells were segmented using InstanSeg^81^ optimized for CRISP output, and per-cell staining intensities were quantified and preprocessed using HFcluster for downstream analysis in Scanpy.

### RNA-seq

Total RNA was extracted using TRIzol and the Direct-zol RNA MiniPrep Plus kit (Zymo Research). RNA integrity was assessed on an Agilent TapeStation. Poly-A-selected RNA-seq libraries were prepared by the SBP Medical Discovery Institute Genomics Core following standard Illumina protocols and sequenced on the Element Biosciences Aviti platform to a depth of approximately 30 million reads per sample.

Raw sequencing reads were quality-assessed using FastQC (v0.11.8), and adaptor sequences were trimmed with Trim Galore (v0.4.4) where necessary. Reads were aligned to the mm10 reference genome using STAR (v2.5.3a), and gene-level counts were generated using HOMER’s analyzeRepeats.pl script. Differential expression analysis was performed using DESeq2 (v1.30.0). Genes with fewer than 10 total raw counts across all samples were excluded prior to normalization. Differentially expressed genes were defined as those with an absolute log2 fold change greater than 1 and an FDR-adjusted p-value < 0.05. GO analysis for RNA-seq was performed using ShinyGO^82^, REVIGO^83^, and Metascape^84^. Upstream regulator analysis was performed using Ingenuity Pathway Analysis (IPA) and filtered to include only transcription factors.

### ATAC-seq

Assay for transposase-accessible chromatin sequencing (ATAC-seq) was performed using the Active Motif ATAC-seq kit. Briefly, 100,000 freshly isolated nuclei were tagmented as previously described^85^, followed by library generation and purification. Libraries were sequenced on the Element Biosciences Aviti platform at the SBP Medical Discovery Institute Genomics Core.

Sequencing reads were trimmed and aligned to the mouse genome (mm10) using Bowtie2, with mitochondrial reads discarded. Multi-mapping and duplicate reads were removed, and alignments were shifted using Deeptools alignmentSieve (v3.4.3). Peaks were called using MACS2 (v2.2.9.1), and reproducible peaks across replicates (n=2) were identified using IDR (v2.0.4.2). Condition-specific peak sets were merged using bedtools (v2.29.2), excluding ENCODE blacklisted regions, yielding 75,164 consensus peaks. Peaks were annotated using HOMER annotatePeaks.pl. The fraction of reads in peaks (FRiP) scores was used to normalize signal tracks. Differential chromatin accessibility was assessed using edgeR (v3.38.4), and profile plots were generated using Deeptools.

### Cut&Tag

Cleavage Under Targets and Tagmentation (CUT&Tag) was performed using the Active Motif CUT&Tag-IT Assay Kit^86^. Nuclei were isolated from mouse liver tissue using Miltenyi gentleMACS C-tubes. For each reaction, 300,000 nuclei were used. CUT&Tag was performed following the manufacturer’s instructions. Library quality and fragment size distribution were assessed on an Agilent TapeStation prior to sequencing. Libraries were sequenced on the Element Biosciences Aviti platform at the SBP Medical Discovery Institute Genomics Core.

Paired-end reads were aligned to the mouse genome (mm10) using Bowtie2. For normalization, reads were separately aligned to the *Escherichia coli* genome. Aligned reads were filtered to remove unmapped reads and converted to fragment BED files. Peaks were called using MACS2 (v2.2.9.1). Normalized signal tracks (bedGraph/bigWig) were generated using the scaling factors. Heatmaps and profile plots were generated using deepTools computeMatrix and plotHeatmap.

### Histology and stainings

Tissues were processed for paraffin embedding and sectioned at 5 μm thickness by the Histology Core Facility at Sanford Burnham Prebys Medical Discovery Institute. Sections were mounted on glass slides, dried at 58°C for 1 h, deparaffinized, and rehydrated through a graded alcohol series. Hematoxylin and eosin (H&E) staining was performed using a Leica Autostainer XL (Leica Microsystems). For collagen detection, Picro-Sirius Red staining was performed on FFPE sections by incubating in Picro-Sirius Red solution (0.1% Sirius Red in saturated picric acid) for 1 h, followed by washes in acidified water, dehydration, and mounting.

### Targeted quantification of cholesterol and cholesteryl esters

Cholesterol (Cho) and cholesteryl esters (ChEs) were quantified by targeted liquid chromatography-mass spectrometry (LC-MS) performed by the Salk Institute Mass Spectrometry Core Facility. Lipids were extracted from liver samples using a modified Bligh-Dyer method. Briefly, samples were homogenized in PBS and extracted with methanol and chloroform containing internal standards (d7-cholesterol). The organic phase was collected, dried under nitrogen, and reconstituted in chloroform:methanol (2:1) for analysis. Samples were analyzed on a Vanquish HPLC coupled to a Q-Exactive quadrupole-orbitrap mass spectrometer equipped with electrospray ionization (Thermo Fisher Scientific). Chromatographic separation was achieved using a C4 column with a gradient of aqueous and isopropanol-based mobile phases. Data were acquired in positive and negative ionization modes using data-dependent MS/MS acquisition. Cho and individual ChE species were identified and quantified by targeted peak integration, with results normalized to internal standards and sample input.

### Blood Chemistry Analysis

Blood was collected via retroorbital bleeding into lithium heparin-coated tubes. Whole blood (100 μL) was loaded into VetScan Mammalian Liver Profile reagent rotors (Zoetis) and analyzed on a VetScan VS2 Chemistry Analyzer within 60 min of collection.

### DNA methylation array

Genomic DNA was extracted using the Monarch Genomic DNA Purification Kit (New England BioLabs). DNA quality and concentration were assessed by TapeStation analysis at the UCSD Institute for Genomic Medicine (IGM) Genomics Core. Samples were processed on the Infinium Mouse Methylation BeadChip (Illumina) according to the manufacturer’s instructions and scanned on an Illumina iScan system.

Methylation array data were processed using SeSAMe (v1.18.4). CpG probes present in only one experimental group were excluded prior to analysis. Differentially methylated loci were defined as those with a Benjamini-Hochberg adjusted p-value < 0.05.

## Data Availability

All datasets generated in this study have been deposited in the Gene Expression Omnibus (GEO) and will be publicly available as of the date of publication.

## Declaration of generative AI

During the preparation of this manuscript the author(s) used Claude (Anthropic) in order to assist with editing and refining manuscript text. After using this tool, the author(s) reviewed and edited the content as needed and take(s) full responsibility for the content of the published article.

## Acknowledgements

Work in the lab of PDA was supported by P01 AG031862 and P01 AG073084. RA and MGT were supported by the California Institute for Regenerative Medicine grant EDUC4-12813. MGT was also supported by an American Society of Hematology (ASH) Hematology Inclusion Pathway (HIP) Graduate Student Award. KM was supported by NIH K99 AG073450. We thank the SBP Bioinformatics Core (Rabi Murad) and the SBP Functional Genomics Core (Chun-Teng Huang) for technical assistance. SBP Core Facilities were supported by NCI CCSG P30 CA030199-44. This work was supported by the Mass Spectrometry Core of the Salk Institute (RRID:SCR_014843) with funding from NIH-NCI CCSG P30 CA014195, NIH-NIA San Diego Nathan Shock Center P30 AG068635, two NIH Shared Instrumentation Grants S10-OD021815 (ThermoFisher Q-Exactive quadrupole orbitrap) and S10-OD038262 (ThermoFisher Orbitrap IQ-X Tribrid), and the Helmsley Center for Genomic Medicine. We thank Kota Kaneko for technical guidance on performing the 70% partial hepatectomy procedure. We thank all members of the Adams lab for critical discussions.

---

**Supplementary Figure 1:**
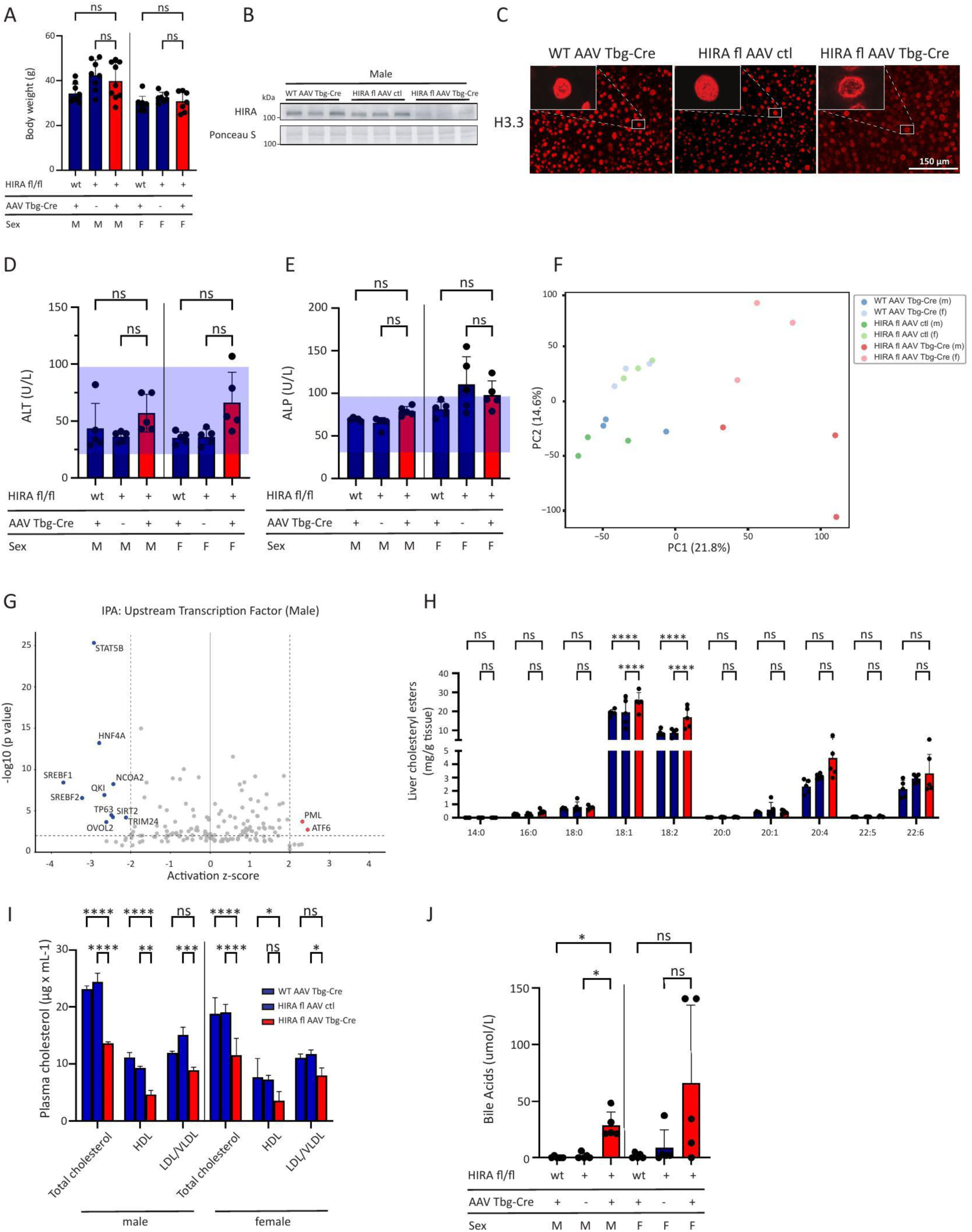
Body weight of control and HIRA KO mice at the time of tissue collection. Male mice: n = 9 (WT AAV Tbg-Cre, HIRA^fl/fl^ AAV Tbg-Cre); n = 8 (HIRA^fl/fl^ AAV ctl), Female mice: n = 9 (WT AAV Tbg-Cre); n = 7 (HIRA^fl/fl^ AAV ctl, HIRA^fl/fl^ AAV Tbg-Cre). B. Representative western blot demonstrating HIRA depletion in liver lysates from control and HIRA KO male mice. Ponceau S staining was used as loading control. n = 8 (HIRA^fl/fl^ AAV ctl); n = 9 (WT AAV Tbg-Cre, HIRA^fl/fl^ AAV Tbg-Cre). C. Immunofluorescence for H3.3 in control and HIRA KO livers, representative of n = 3 experiments. Scale bar, 150 μm. D. Effect in control and HIRA KO mice on serum ALT levels. n = 5 per group. E. Effect in control and HIRA KO mice on serum ALP levels. n = 5 per group. F. RNA-seq principal component analysis. n = 3 per group. G. Ingenuity Pathway Analysis (IPA) showing top predicted upstream transcription factors regulating DEGs in control versus HIRA KO mice. H. Hepatic cholesteryl ester composition showing individual species (14:0-22:6) as determined by LC-MS analysis. n = 5 per group. I. Plasma lipid profile showing total cholesterol and lipoprotein fractions (HDL, and LDL/ VLDL) in control and HIRA KO mice. n = 3 per group. J. Serum bile acid levels in control and HIRA KO mice measured using VetScan VS2 Mammalian Liver Profile rotors. n = 5 per group. Data are presented as means ± SEM. Statistical significance was determined by one-way analysis of variance (ANOVA) with Tukey’s multiple comparisons test. Asterisk (*) indicates p < 0.05, (**) p < 0.01, (***) p< 0.001, (****) p < 0.0001.

**Supplementary Figure 2:**
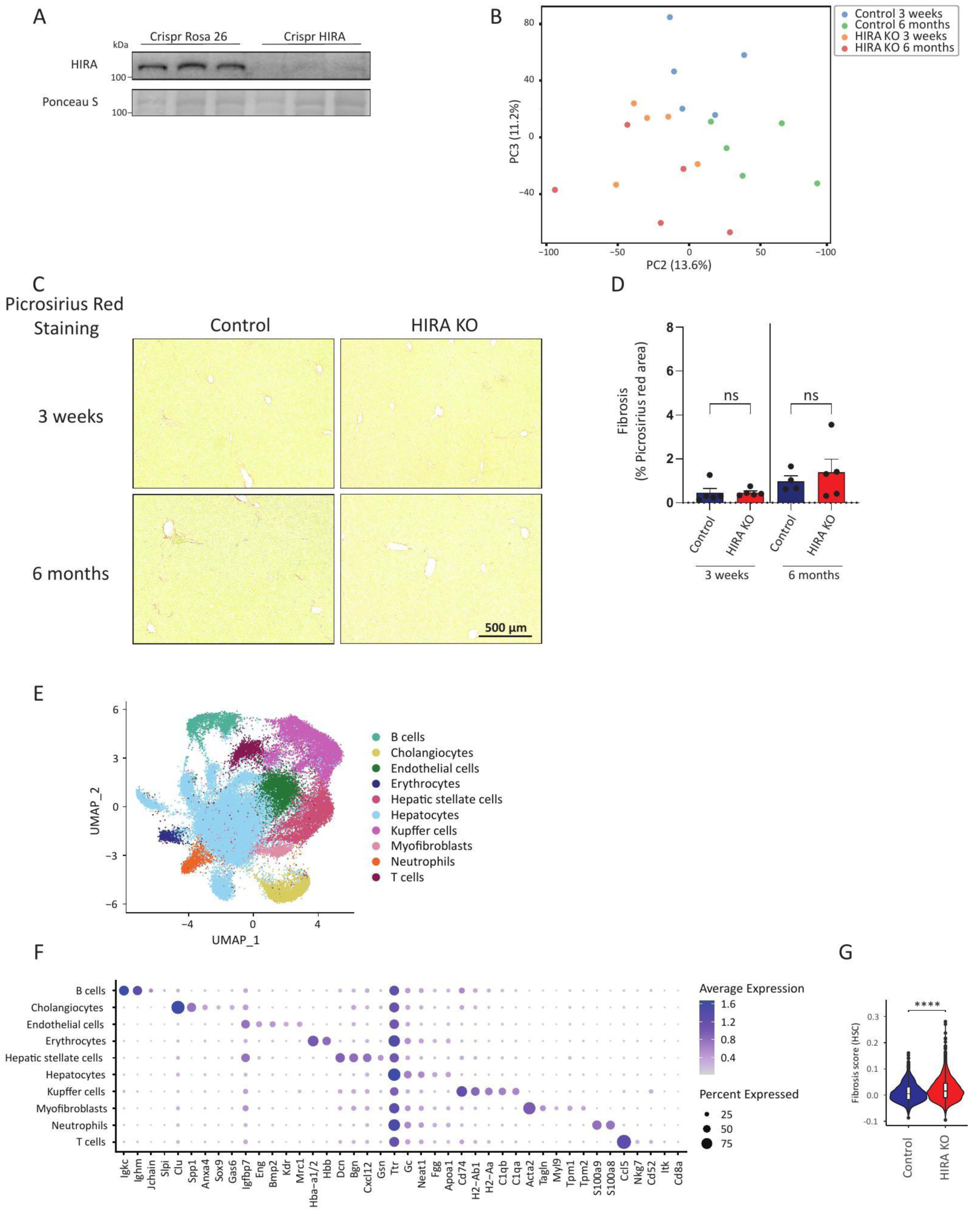
A. Representative western blot showing HIRA depletion in liver lysates from control and HIRA KO mice generated using CRISPR-Cas9. Ponceau S staining was used as loading control. n = 5. B. RNA-seq principal component analysis. n = 5 per group. C. Representative picrosirius red stainings of liver sections from control and HIRA KO mice at 3-week and 6-month timepoints. Scale bar, 500 μm. D. Quantification of picrosirius red positive area. n = 5. E. UMAP visualization of cell populations identified by CosMX spatial transcriptomics analysis in control and HIRA KO mouse liver tissue. n = 3. F. Dot plot displaying the top 5 marker genes for each cell type identified by CosMx spatial transcriptomics. Dot size represents the percentage of cells expressing each gene; color intensity indicates average expression level. G. Fibrosis score derived from a 43-gene signature (Suppl. Table 1) encompassing extracellular matrix components, matrix remodeling enzymes, and TGF-β signaling genes in control and HIRA KO liver sections. n = 3 per group. Data are presented as means ± SEM. Statistical significance for D was determined by one-way analysis of variance (ANOVA) with Tukey’s multiple comparisons test. For G, the Wilcoxon rank-sum test was used. Asterisk (*) indicates p < 0.05, (**) p < 0.01, (***) p< 0.001, (****) p < 0.0001.

**Supplementary Figure 3:**
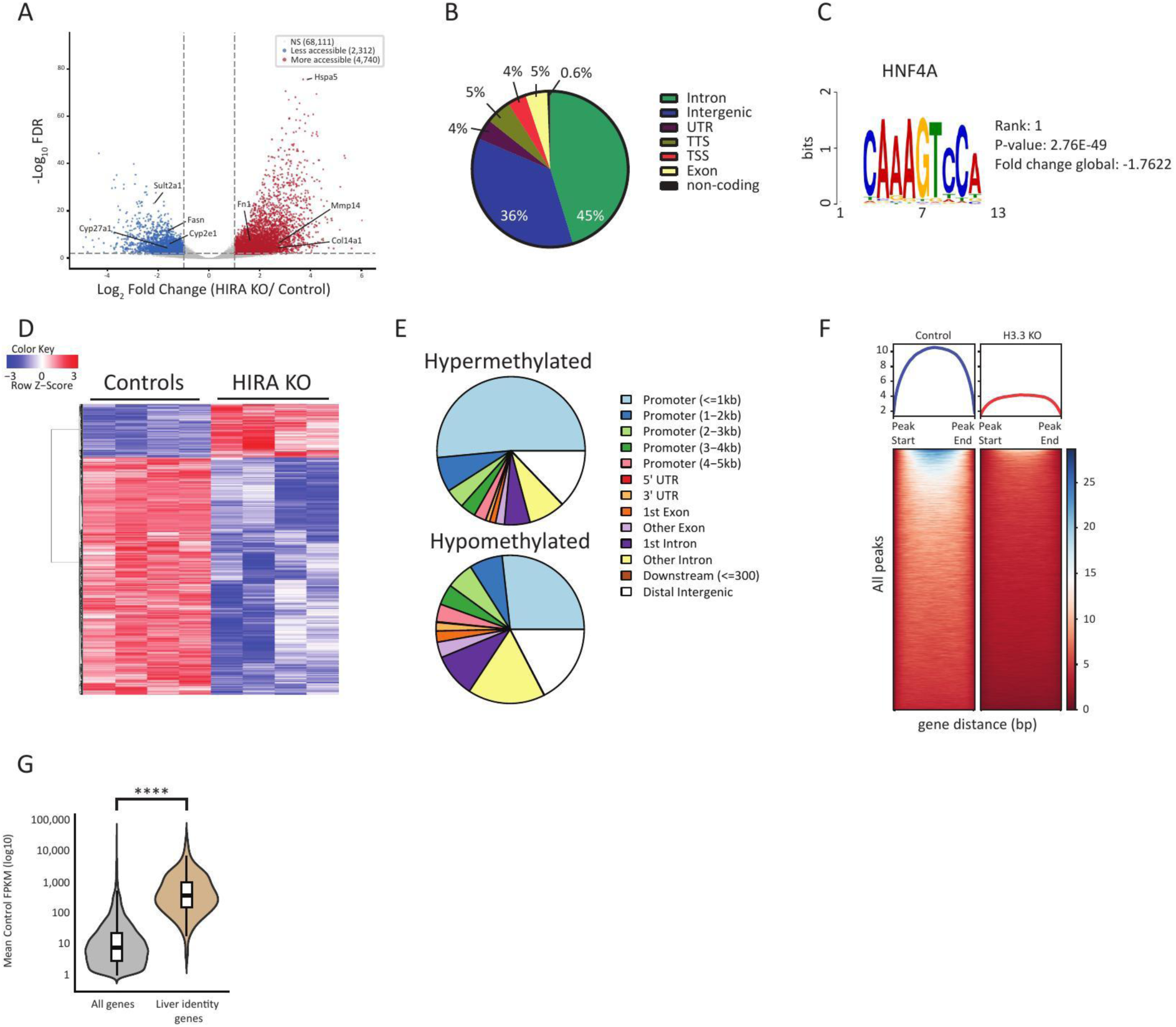
A. Volcano plot of differentially accessible chromatin regions identified by ATAC-seq in HIRA KO versus control livers. Regions classified as less accessible (n = 2,312), more accessible (n = 4,740), and not significant (NS; n = 68,111) are indicated. Selected genes are labeled. n = 2 per group. B. Pie chart showing the genomic distribution of differentially accessible ATAC-seq peaks by functional annotation (intron, intergenic, UTR, TTS, TSS, exon, non-coding). C. Top enriched transcription factor motif at regions with decreased chromatin accessibility in HIRA KO livers identified by ATAC-seq. HNF4α was ranked first. D. Heatmap of differentially methylated CpG sites (35,594 hypomethylated, 7,968 hypermethylated) in control and HIRA KO livers. n = 4 per group. E. Genomic distribution of hypermethylated and hypomethylated CpG sites annotated by functional category. F. Heatmap of H3.3 CUT&Tag signal intensity across all identified peaks in control and H3.3 KO livers, validating the H3.3 CUT&Tag approach. G. Expression distribution of all expressed genes compared to liver identity genes in female mice. Wilcoxon rank-sum test (Mann-Whitney U test) was used to calculate statistical significance. Asterisk (*) indicates p < 0.05, (**) p < 0.01, (***) p< 0.001, (****) p < 0.0001.

**Supplementary Figure 4:**
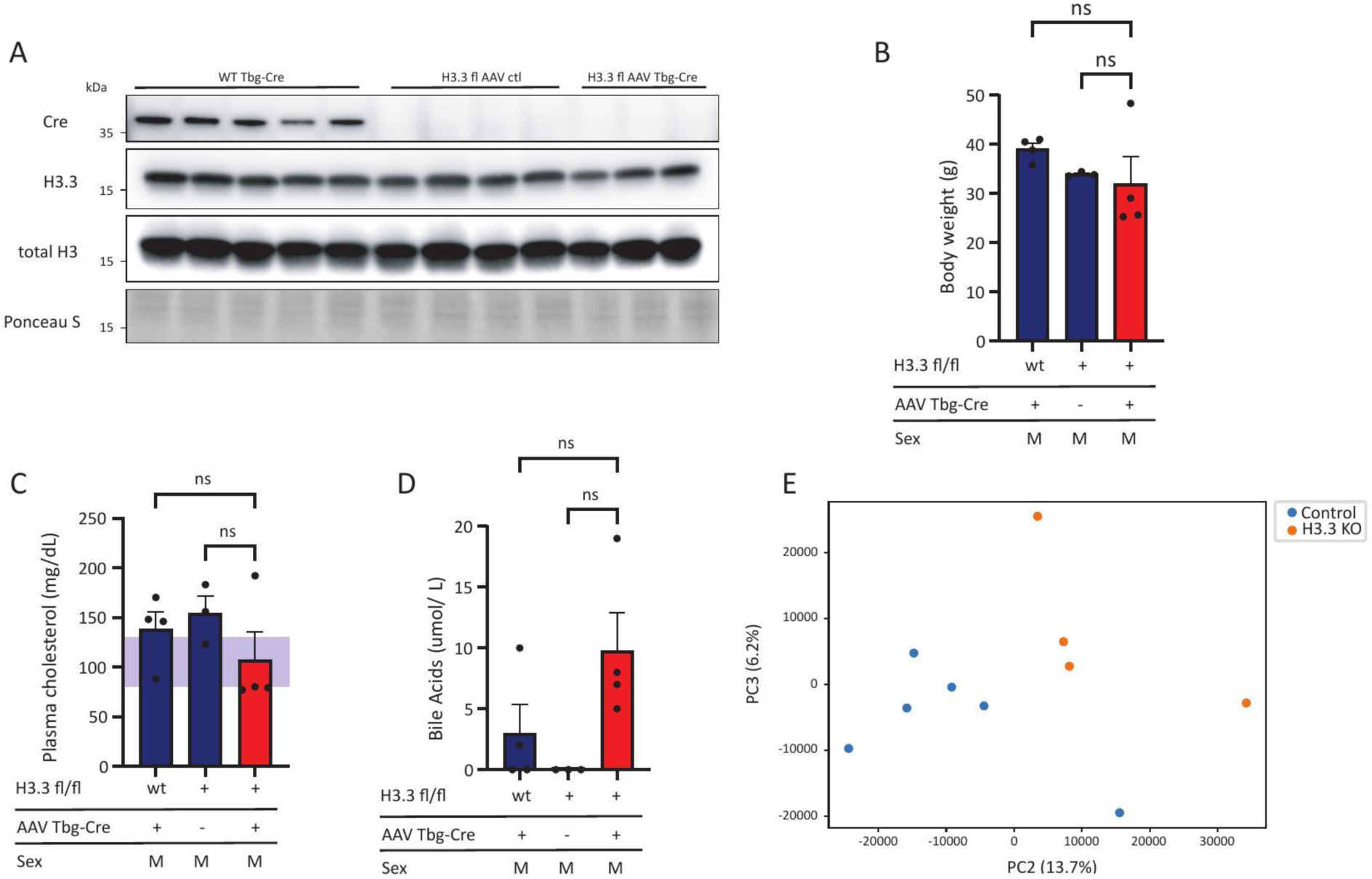
A. Representative western blot showing Cre, H3.3 and total H3 protein levels in liver lysates from control and H3.3 KO mice 1 year post-knockout. Ponceau S staining served as loading control. n = 5 (WT AAV Tbg-Cre); n = 4 (H3f3a^fl/fl^ H3f3b^fl/fl^ AAV ctl); n = 3 (H3f3a^fl/fl^ H3f3b^fl/fl^ AAV Tbg-Cre). B. Body weight of control and H3.3 KO mice 3 weeks post-knockout at the time of tissue collection. n = 4 (WT AAV Tbg-Cre, H3f3a^fl/fl^ H3f3b^fl/fl^ AAV Tbg-Cre); n = 3 (H3f3a^fl/fl^ H3f3b^fl/fl^ AAV ctl). C. Plasma cholesterol levels in control and H3.3 KO mice. D. Serum bile acid levels in control and H3.3 KO mice measured using VetScan VS2 Mammalian Liver Profile rotors. E. RNA-seq principal component analysis. n = 3 (WT AAV Tbg-Cre, H3f3a^fl/fl^ H3f3b^fl/fl^ AAV ctl); n = 4 (H3f3a^fl/fl^ H3f3b^fl/fl^ AAV Tbg-Cre). Data are presented as means ± SEM. Statistical significance was determined by one-way analysis of variance (ANOVA) with Tukey’s multiple comparisons test. Asterisk (*) indicates p < 0.05, (**) p < 0.01, (***) p< 0.001, (****) p < 0.0001.

**Supplementary Figure 5:**
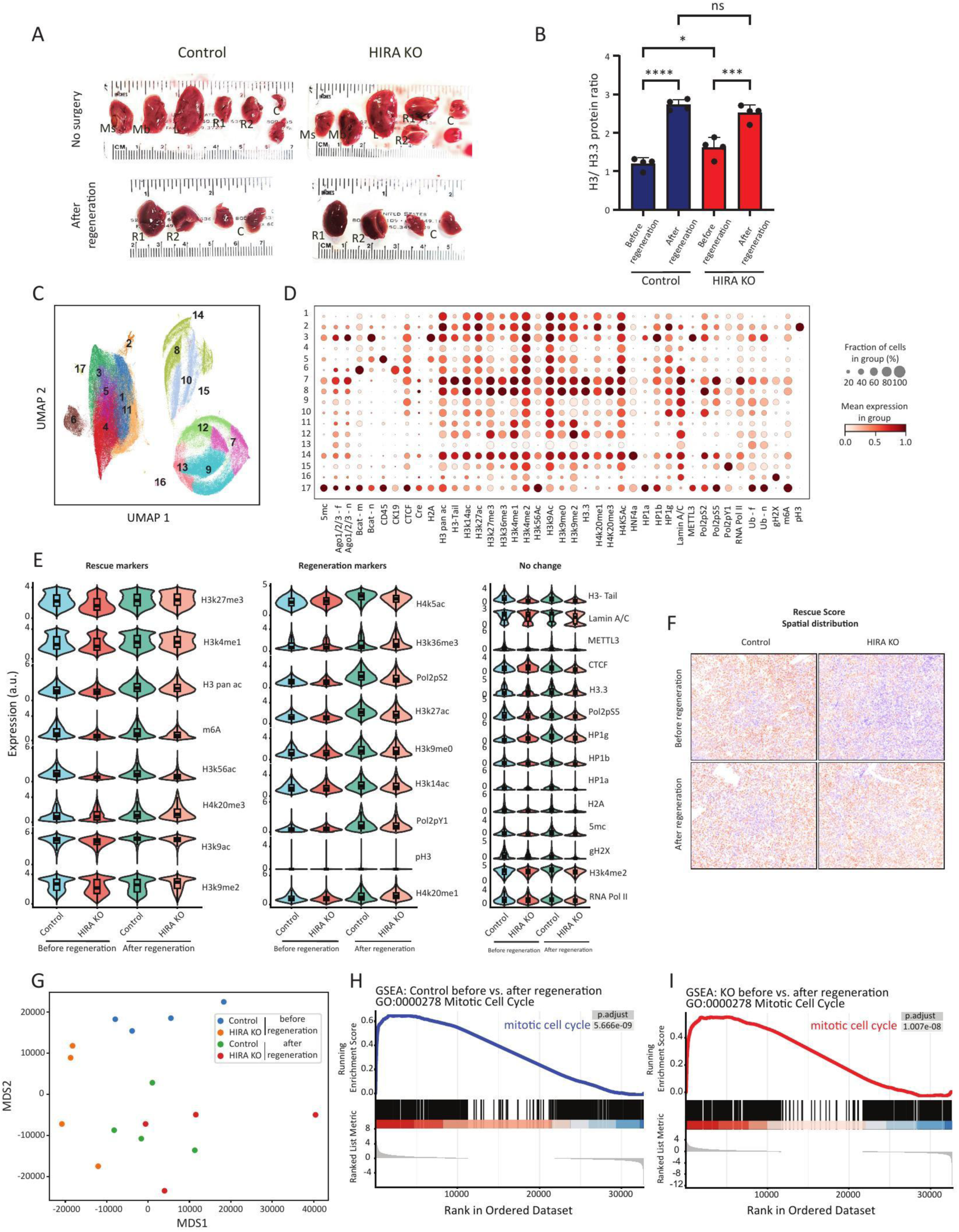
A. Representative images of liver lobes from control and HIRA KO mice without surgery and after regeneration. R1, R2: right lobe segments; C: caudate lobe; Ms: median lobe segments; L: left lobe. B. Western blot quantification, calculating the ratio of total H3/ H3.3 in control and HIRA KO livers before and after regeneration. n = 4. C. UMAP visualization identified by CODEX multiplex immunofluorescence analysis in control and HIRA KO liver tissue before and after regeneration. D. Dot plot displaying expression of epigenetic markers and cell type markers across identified clusters. Dot size represents the fraction of cells expressing each marker; color intensity indicates mean expression level. E. Violin plots showing expression of individual epigenetic markers in hepatocytes (HNF4α^+^, CK19^−^, CD45^−^; n = 194 165 cells) across conditions. Markers are grouped into rescue markers (altered by HIRA loss and restored after regeneration), regeneration markers (changed with regeneration in both genotypes), and no change markers (largely stable across conditions). F. Spatial distribution of rescue score mapped onto tissue sections from control and HIRA KO livers before and after regeneration. G. RNA-seq principal component analysis (MDS plot) of control and HIRA KO liver samples before and after regeneration. n = 4 per group. H, I. Gene Set Enrichment Analysis (GSEA) showing enrichment of mitotic cell cycle genes (GO:0000278) in control (H) and HIRA KO (I) livers comparing before versus after regeneration. Data are presented as means ± SEM. Statistical significance was determined by two-way ANOVA with Tukey’s multiple comparisons test. Asterisk (*) indicates p < 0.05, (**) p < 0.01, (***) p< 0.001, (****) p < 0.0001.

